# A variant rRNA serves as a translational repressor in *Plasmodium falciparum*

**DOI:** 10.64898/2026.06.17.732804

**Authors:** Tiziano Vignolini, Olivia Carril, Victor Tobiasson, Joe Georgeson, Miguel A. Garcia-Campos, Justine E. Couble, Grégory Doré, Donna Matzov, Sebastian Hutchinson, Jessica M. Bryant, Moran Shalev-Benami, Schraga Schwartz, Sebastian Baumgarten

## Abstract

Ribosome composition can vary through differences in associated proteins, post-transcriptional and post-translational modifications. Such heterogeneity enables ribosomes to respond to environmental^1^ or pathological^2,3^ conditions, and modulate localized translation^4^. A long-standing observation has also been the differential expression of variant ribosomal RNA (rRNA) alleles across developmental^5–7^ or cellular states^8–14^. Yet how exchanging the catalytic ribosome core could regulate translational outcomes remains unknown. Here, we report the functional characterization of a genomically-encoded, divergent rRNA that serves as a dominant-negative repressor of translation during host-to-vector transmission in the human malaria parasite. This allele only encodes for large subunit rRNAs, lacks ITS2 splicing, yet retains conserved rRNA modification and folding patterns alongside vast expansion segments. The resulting large subunit engages mRNA at translation start sites but appears to elongate inefficiently, likely due to divergences in the peptidyl transferase center obstructing the exit tunnel. Through its precisely timed transcription immediately after transmission, this rRNA represses mRNAs that were highly translated in the human, facilitating the transition of the translational program for mosquito-stage development. Our data identify a repressive ribosome population whose antagonistic function is encoded by an independently evolved, variant rRNA allele, defining the conceptual foundation for an additional layer of inherent translational regulation.

## RESULTS

Ribosomes are the universally conserved, multi-component macromolecular complexes comprised of a catalytic, ribosomal RNA (rRNA) core and multiple associated proteins that are responsible for protein synthesis in all living organisms. While all known ribosomes translate mRNA into proteins, their composition regarding the association with ribosomal proteins^2,15^ or post-transcriptional modifications^1,3^ can vary within the same organism between cell types^16^, different subcellular localizations^4^ or in response to external stimuli^1,2^. Such compositional heterogeneity around the same rRNA core can provide a capacity to fine tune translational outputs across different cellular states^17^. rRNA itself is encoded by multiple repeats of rRNA alleles from which 18S (making up the small subunit, SSU), 5.8S and 28S (comprising the large subunit, LSU) is transcribed (Fig. 1a). For many years, it has been observed that not all rRNAs are equally transcribed. Instead, distinct rRNA alleles with variable sequences can be differentially expressed during cell development^5–7^, in various human disease states^8,9^ or during exposure to stress in bacteria, where they can facilitate a translational response^14,18^. Yet how specific variations in the rRNA sequence can provide an inherent, regulatory role to a distinct ribosome population remains unclear.

**Figure 1.**
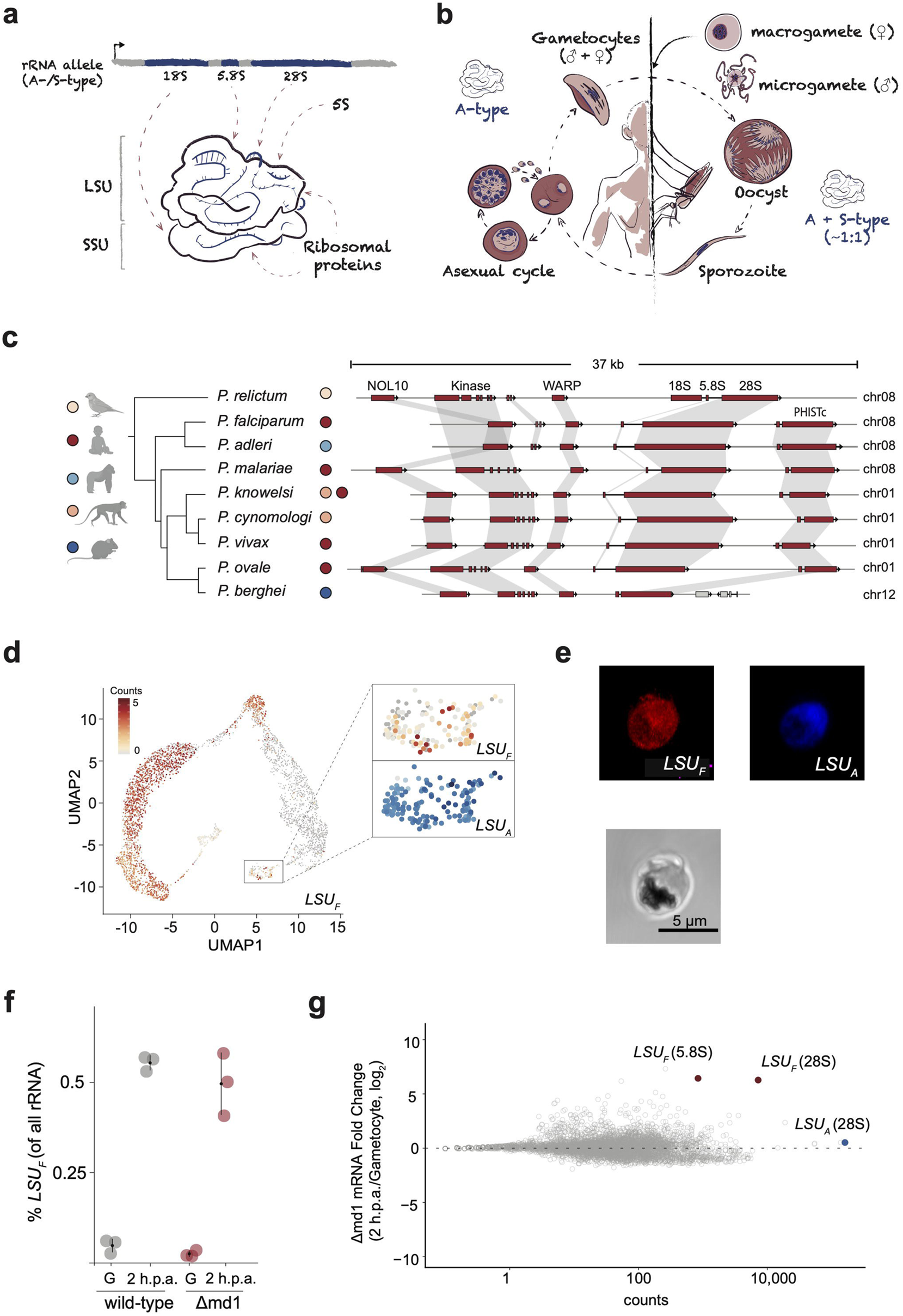
*LSU_F_* rRNA is a marker of *P. falciparum* transmission. (a) Schematic of the composition of a eukaryotic ribosome. 18S, 5.8S and 28S are transcribed as one polycistronic transcript. 18S assembles into the small subunit (SSU), whereas 5.8S and 28S form the catalytic core of the large subunit (SSU). The 5S rRNA is transcribed from a separate locus and is incorporated in the LSU. The arrow on the rRNA allele indicates the direction of transcription. (b) Diagram of the *P. falciparum* lifecycle including the developmental transition from gametocytes to female (left) and ‘exflagellating’ male (right) gametes during host-to-vector transmission. (c) Synteny of the F-type rDNA locus across *Plasmodium* species. (d) scRNA-seq expression profile of *P. falciparum LSU_F_* 28S during development in the *An. gambiae* mosquito^20,25^. Developmental stages are shown in Supplementary Fig. 1d. (e) RNA-FISH of *LSU_F_* (red) and *LSU_A_* (blue) 28S rRNA in gametes at 2 h.p.a. Bottom panel: Brightfield image of the gamete. Scale bar: 5 µm. (f) Percentage of *LSU_F_*28S transcripts among all LSU rRNA (i.e. *LSU_F_* + *LSU_A_*) measured by RT-qPCR in mature gametocytes (G) and gametes (2 h.p.a.) of wild-type (WT) and a male-deficient, female-only cell line (Δmd1)^34^. (g) Changes of transcript abundance between Δmd1 gametocytes and gametes at 2 h.p.a. calculated from mRNA-seq. *LSU_F_*rRNA is shown in red, *LSU_A_* 28S rRNA in blue.

One of the most compelling instances of such rRNA-mediated ribosome heterogeneity is *Plasmodium falciparum*, a unicellular, eukaryotic parasite and causative agent of the most virulent form of human malaria. Instead of encoding multiple rRNA repeats that together comprise the nucleolar organizer region in most eukaryotes^19^, *P. falciparum* contains only five canonical rRNAs (i.e. encoding a 18S, 5.8S and 28S) that are each located on a different chromosome. On top, the parasite tightly controls the expression of each of the five rRNA alleles as it progresses through the human host and *Anopheles* vector (Fig. 1a,b)^12^; ‘A-type’ rRNA are expressed throughout the parasite lifecycle, whereas ‘S-type’ rRNA transcription is restricted to the developmental stages in the vector (Fig. 1b)^12,20^, leading to heterogeneous ribosome populations during the parasite’s developmental journey through the mosquito^12^. This pattern of variant rRNA transcription makes *P. falciparum* an evolutionary ‘extreme’ that could highlight previously unseen avenues to decipher a possibly rRNA-mediated, regulatory function of eukaryotic ribosomes. However, A- and S-type rRNA feature high sequence similarity that largely impedes genomic analyses, and only very low amounts of biological material can be retrieved from mosquito stage parasites. These limitations have so far restricted studies on the different rRNA variants to phenotypic descriptions^21,22^ and precluded mechanistic insight into distinct roles of different ribosome populations. Thus, whether variant rRNA alleles could provide a genomically-encoded, regulatory capacity for ribosomes remains to be discovered in any eukaryote.

### A divergent rRNA allele is a marker of *P. falciparum* transmission

In addition to the two known classes of rRNA, we recently detected the expression of an additional, highly divergent rRNA allele during parasite development in the mosquito midgut (i.e. during the ‘oocyst’ stage, eight days after *Anopheles* infection)^12^. The corresponding rDNA locus lacks an 18S SSU rRNA, but encodes an expanded and highly divergent 28S rRNA that is 50% larger (6,174 nt) and shares only 50% sequence identity with any of the canonical 28S sequences encoded by the parasite. Correspondingly, the original genome annotation of *P. falciparum* annotated this locus (5.8S: PF3D7_0801200, 28S: PF3D7_0801100) as a ‘fragment rRNA’^23^, and was therefore named ‘F-type’ large subunit (*LSU_F_*) rRNA. Searching available *Plasmodium* genomes, we find that *LSU_F_* rDNA with a similar, incomplete layout are conserved and located at syntenic genomic loci in all primate-infecting *Plasmodium* species but is lost in rodent-infecting malaria parasites (Fig. 1c, Supplementary Fig 1a). In contrast, the rDNA gene that is located at the corresponding genomic position in the early-branching, bird-infecting *P. relictum* genome encodes a canonical rRNA, suggesting that *LSU_F_*independently evolved from a canonical rRNA allele after the switch to mammalian hosts^24^ (Fig. 1c).

*LSU_F_* rRNA is not detectable in *P. falciparum* parasites that asexually replicate in the human host, and during development in the mosquito midgut makes up < 1% of the entire LSU rRNA content^12^. In contrast to other transcriptionally silent rDNA loci (namely ‘S-type’), the *LSU_F_*rDNA locus is not subject to heterochromatinization during asexual proliferation in the host^12^. Instead, it exhibits an accessible upstream chromatin configuration, suggesting that *LSU_F_*rDNA is in a transcriptionally permissive state and may support rapid transcriptional activation (Supplementary Fig. 1b). Profiling available expression data of *LSU_F_* homologs revealed its expression also during *P. malariae* midgut development, and high transcript abundance of *LSU_F_*were also detectable during liver development in *P. cynomologi* and *P. vivax* (Supplementary Fig. 1c). Reanalyzing recent gene expression data of *P. falciparum* similarly showed that *LSU_F_* is expressed in the hepatic stages, reaching levels similar to those measured during development in the mosquito (Supplementary Fig. 1c). Resolving *LSU_F_* transcript abundance across *P. falciparum* development in the mosquito even further using scRNA-seq^20^ confirmed its expression in the oocyst stage, starting ∼2 days after mosquito infection. However, this data also revealed a second time of expression immediately after *P. falciparum* is transmitted from the human to the mosquito (Fig. 1d)^25^.

To successfully transmit to a mosquito vector, haploid *P. falciparum* parasites that asexually replicate within human red blood cells develop into non-replicative, male and female transmission stages^26^ (Fig. 1b). Following uptake by an *Anopheles* during a bloodmeal, these so-called gametocytes rapidly differentiate within minutes. Female gametocytes develop into a single macrogamete, whereas male gametocytes undergo three rounds of genome replication, followed by the ‘exflagellation’ of the resulting eight motile microgametes.^27^ Inside the mosquito gut, these gametes fuse to a zygote which develops into a motile ookinete that exits the gut. Attached to the outside of the mosquito midgut, multiple rounds of asexual replication within ‘oocysts’ lead to the development of sporozoites. These human-infective stages are transmitted to a new host during the next bloodmeal, where they first replicate within liver cells before infecting red blood cells again (Fig. 1b)^20^. Importantly, for female parasites the transition from a semi-quiescent gametocyte in the human to a gamete inside the mosquito gut (also called gametocyte ‘activation’) is regulated almost exclusively on the translational level. These gametocytes already transcribe the mRNAs that encode for proteins needed in the mosquito, but repress their translation using a dedicated complex assembled around the DDX6 helicase ‘DOZI’ until they are released after transmission^28–31^. *P. falciparum* gametes are readily available from *in vitro* cultures without the need of *Anopheles* mosquitoes^32,33^, allowing the application of a range of multi-omics approaches.

A direct comparison with ‘A-type’ large subunit (*LSU_A_*) rRNA expression and fluorescent in situ hybridization (RNA-FISH) shows that the two rRNA types are simultaneously present within the same female macrogamete (Fig. 1d,e Supplementary Fig. 1d). Comparisons of *LSU_F_* rRNA expression in wild-type parasites during *in vitro* gametocytes to gamete development (i.e. 2 hours post activation [h.p.a.]) recapitulates the upregulation observed by single-cell RNA-seq, with *LSU_F_* making up ∼0.5% of all LSU rRNA, the canonical *LSU_A_* rRNA comprising the remaining rRNA pool (Fig. 1f). A similar upregulation is observed for a male deficient strain (Δmd1)^34^, suggesting that *LSU_F_* rRNA transcription appears to be activated in female gametes and does not depend on fertilization or zygote formation (Fig. 1f). A transcriptome-wide comparison of changes in RNA abundance between female gametocytes and gametes further shows that *LSU_F_* rRNA is among the most upregulated and most abundant transcripts in *P. falciparum* gametes (Fig. 1g, Supplementary Fig. 1e,f, Supplementary Table 1). In contrast, no annotated SSU or LSU ribosomal protein shows a similar upregulation (Supplementary Fig. 1g). Interestingly, while *LSU_F_* rRNA is not detectable in gametocytes (Supplementary Fig. 1f), it is already one of the most upregulated RNA transcripts as soon as 5 min after gametocyte activation (Supplementary Fig. 1h)^30^, identifying *LSU_F_*rRNA as a marker of gametocyte-to-gamete developmental transition.

### *LSU_F_* rRNA encodes a divergent large subunit paralog

To elucidate if *LSU_F_* rRNA retains central characteristics of eukaryotic rRNA, we first performed nanopore-based, direct RNA sequencing (DRS) of gamete total RNA sampled at 2 h.p.a. The canonical ‘A-type’ rDNA locus shows a classical pattern of high read coverage on the 18S, 5.8S and 28S sequences, indicative of efficient splicing of internal transcribed spacer (ITS) 1 and 2 during ribosome biogenesis (Fig. 2a). In contrast, the ITS2 sequence of *LSU_F_* rRNA is not only more than 6 times longer (1,319 nt) than the ‘A-type’ ITS2 (199 nt), but also appears not to be spliced from the 5.8S and 28S sequences (Fig. 2a). High coverage of *LSU_F_* ITS2 is similarly detectable by orthogonal sequencing approaches in 2 h.p.a gametes, oocysts and during hepatocyte development (Supplementary Fig. 2a), as well as in *LSU_F_*homologs (Supplementary Fig. 2b). This indicates that a *LSU_F_* pre-rRNA is not post-transcriptionally cleaved but remains as one contiguous, large subunit rRNA that maintains an ITS2 ‘foot’^35^.

**Figure 2.**
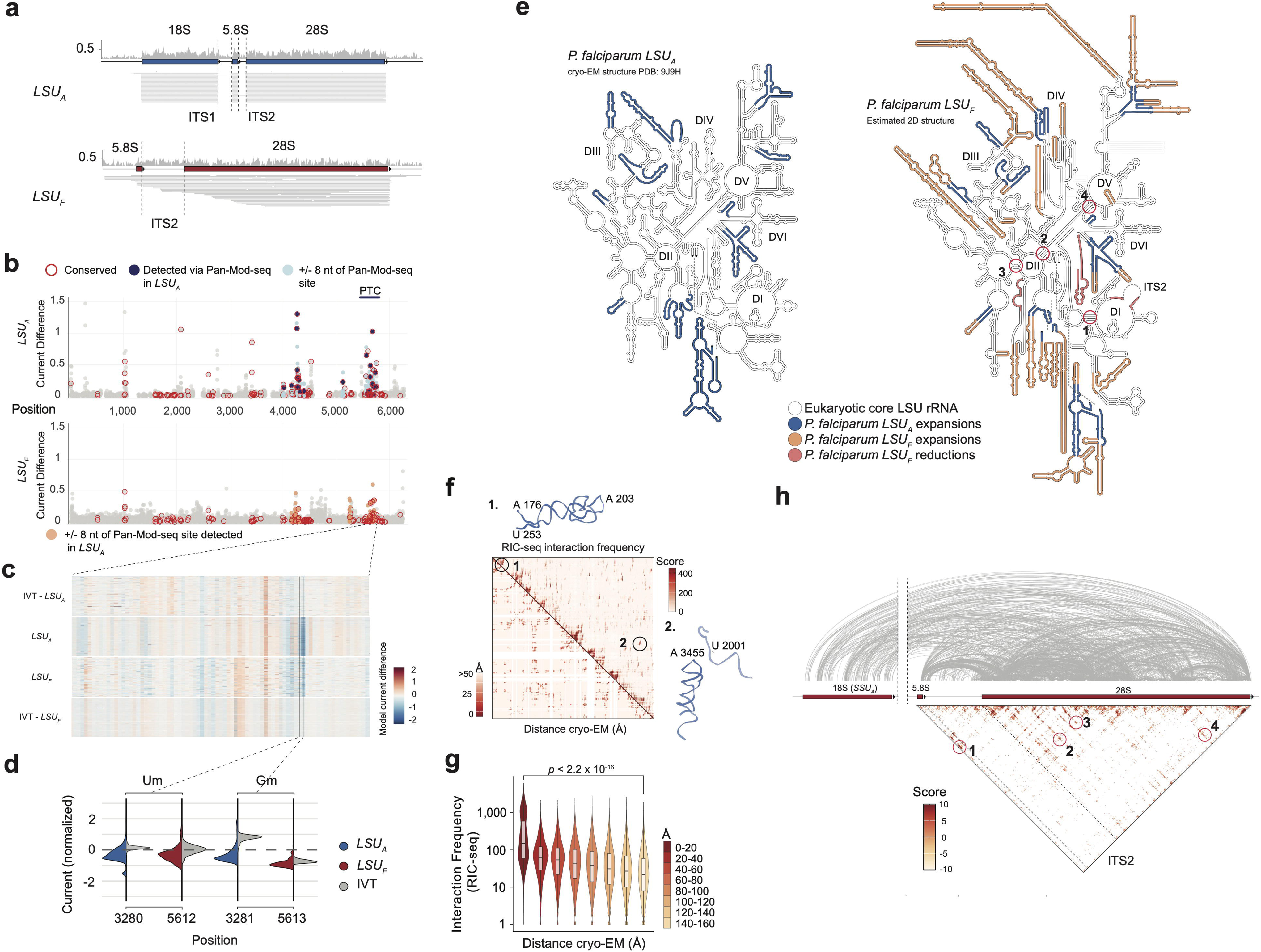
*LSU_F_* rRNA encodes for a divergent large subunit. (a) Alignment of reads generated by direct RNA sequencing to the variant *LSU_F_* rRNA (top) and canonical *LSU_A_* rRNA allele (bottom). GC content (20nt bins) is indicated on the top. (b) Differences of the normalized mean signal current between native and control RNA (IVT) measured by DRS for each nucleotide position in *LSU_A_*(top) and *LSU_F_* (bottom) 28S sequences at 2 h.p.a. (ordinate). The abscissa indicates the nucleotide position from a pairwise alignment between the *LSU_F_* and *LSU_A_* 28S sequences. Red circles indicate positions with evolutionary conserved rRNA modifications (see Extended Data Table 2). Blue sites are detected via Pan-Mod-seq. Light blue sites are located ± 8 nucleotides surrounding a modification that was identified via Pan-Mod-seq in *LSU_A_*, with orange sites highlighting the corresponding positions in *LSU_F_*. (c) Heatmap of read alignments colored by the difference of the observed and expected normalized current level in the PTC of *LSU_F_* and *LSU_A_*28S. The IVT controls are shown above (for *LSU_A_*) and below (for *LSU_F_*). Each row represents one read, with columns denoting the different positions. (d) Distribution of normalized signal currents in native and IVT RNA at two positions that contain a conserved rRNA modification in *LSU_A_* 28S (annotated via Pan-Mod-seq) and the currents in the corresponding position of the *LSU_F_* sequence. (e) Secondary structure map of the canonical *LSU_A_* (left) and *LSU_F_* (right) rRNA. Numbers denote regions of exemplary interactions detected by RIC-seq (see Fig. 2h). Roman numbers denote domains (D). (f) Top right triangle: Balanced contact frequencies (‘Score) representing intramolecular interactions within the *LSU_A_* 28S rRNA identified via RIC-seq. Linear scale, 5 nt resolution. Bottom left triangle: Heatmap representing spatial distances (in Å) of the 28S within *LSU_A_* obtained via cryo-EM (PDB: 9J9H) at 5 nt resolution. Darker shades of red represent increased contact frequency and closer spatial distance for the RIC-seq and cryo-EM model, respectively. Numbers in the heatmap highlight cryo-EM interactions detected by RIC-seq shown above and on the right. (g) Comparison of binned spatial distance (abscissa, measured via cryo-EM) and contact frequency (ordinate, measured via RIC-seq) within the *LSU_A_* 28S rRNA. *p-*value was calculated using an unpaired Wilcoxon Rank Sum test. (h) Intermolecular interactions between the *LSU_F_* rRNA and the S*SU_A_* 18S rRNA (top). Balanced contact frequencies (Score) of intramolecular interactions within the *LSU_F_* rRNA. Numbers denote exemplary interactions shown in the secondary structure of Fig. 2e. Linear scale, resolution: 10 nt.

rRNAs are highly modified, with multiple positions being conserved across pro- and eukaryotes^36,37^. Mapping these modifications across *P. falciparum* is challenging, due to the high AT content of the genome^23^ and high degree of sequence overlap between different *LSU* variants^12^. Nonetheless, by integrating Pan-Mod-seq^38^, and an evolutionary analysis where we prioritized positions known to be modified in other eukaryotic species, we conservatively identified 27 modifications in the 28S sequence of the canonical *LSU_A_* rRNA from total RNA sampled at 2 h.p.a (Fig. 2b, Supplementary Table 2). All of those modifications were similarly identified using orthogonal direct RNA sequencing (DRS) via the comparison of the mean normalized read current measured in native total RNA and an *in vitro* transcribed (IVT) control RNA (Fig. 2b, Supplementary Fig. 2c, Supplementary Table 2). Overlaying the DRS signal of modification detected in *LSU_A_* with the DRS signals of *LSU_F_*showed that 13 of them are equally detectable in *LSU_F_*, (Fig. 2b), most of them located close to the peptidyl transferase center (PTC, Fig. 2c,d).

In virtually all organisms, LSU rRNA folds into a conserved secondary and tertiary structure underlying the catalytic activity of the ribosome. To understand the divergence of the *LSU_F_* rRNA, we next modeled its secondary structure using a template-based approach^39,40^. The resulting secondary structure mapping revealed a rRNA core that is largely conserved between *LSU_A_* and *LSU_F_* rRNA (Fig. 2e, Supplementary Fig. 2d), yet local regions exhibiting low sequence complexity prevented us from modeling the central protuberance and regions of domain IV. Most strikingly though, *LSU_F_* is decorated with multiple extensive rRNA expansion segments (Fig. 2e). While the secondary structure of the rRNA core is well conserved across all domains of life, many eukaryotes feature such additional rRNA arranged in signature expansion segments. These expansions extend ancestral bacterial helices, and are broadly conserved across eukaryotic lineages (Supplementary Fig. 2e). However, the expansions observed in *LSU_F_* rRNA are distinct due to their prodigious size (11 – 301 nt, Supplementary Table 3) and evolutionary scope, not being shared with *LSU_A_* rRNA, nor with any other eukaryotes^41–43^ (Supplementary Table 3, Supplementary Fig. 2d,e). While *LSU_F_* rRNA decidedly originates within the *Plasmodium* lineage, the nucleotide composition of rRNA tentacles is reminiscent of those in metazoan rRNAs. In both cases, these large linear rRNA expansions have low sequence complexity, exhibit a biased compositions including the degree of polarization, rapid rates of accretion and prominent, repetitive units^41,43^ (Supplementary Fig. 2f). While metazoan expansion segments are GC-biased^44^, the substantial AU-bias in *LSU_F_*mirrors the general AT-bias of intergenic regions in *P. falciparum*^23^ (Supplementary Fig. 2f). In addition, while the overall divergence of *LSU_F_* rRNA is dominated by expansions, prominent helices are reduced or lacking in secondary structure. Specifically, helices 99 and 100 of domain VI as well as helices 9, 10 and 11 flanking the retained ITS sequence are either highly diverged or lost altogether (Fig. 2e, Supplementary Fig. 2d). These deletions, together with a highly unusual internal expansion inside helix 18 likely significantly remodel what would be the membrane facing interface of *LSU_F_*.

To provide experimental evidence for the folding of *LSU_F_* rRNA, we characterized the *in vivo* 3D RNA architecture of *LSU_A_* and *LSU_F_* rRNA using RNA *in situ* conformation sequencing (Supplementary Fig. 3a, ‘RIC-seq’^45^). Analogous to approaches revealing 3D genome architecture, here RNA-protein complexes are first crosslinked, RNA transcripts are partially digested and free RNA ends in close proximity are ligated together (Supplementary Fig. 3b-d). Sequencing of the resulting chimeric RNA molecules can then provide information on the interaction and spatial organization within or between different transcripts in a native cellular context^46^. To confirm the sensitivity of this approach, we first compared intramolecular interaction frequencies of the canonical *LSU_A_* 28S rRNA with spatial distances retrieved from an atomic model generated by cryo-electron microscopy^47^. As expected, we find that interaction frequencies mirror the absolute distances within the LSU, with regions in close spatial proximity featuring higher interaction frequencies than more distal ones (Fig. 2f,g). In addition, we find that *LSU_A_* 28S rRNA has the highest frequency of intermolecular interactions with *LSU_A_* 5.8S and the *SSU_A_* 18S (Supplementary Fig. Fig. 3e) rRNA, thus confirming that this approach can recapitulate distances between spatially associated transcripts and RNA architecture even within a single macromolecular complex.

**Figure 3.**
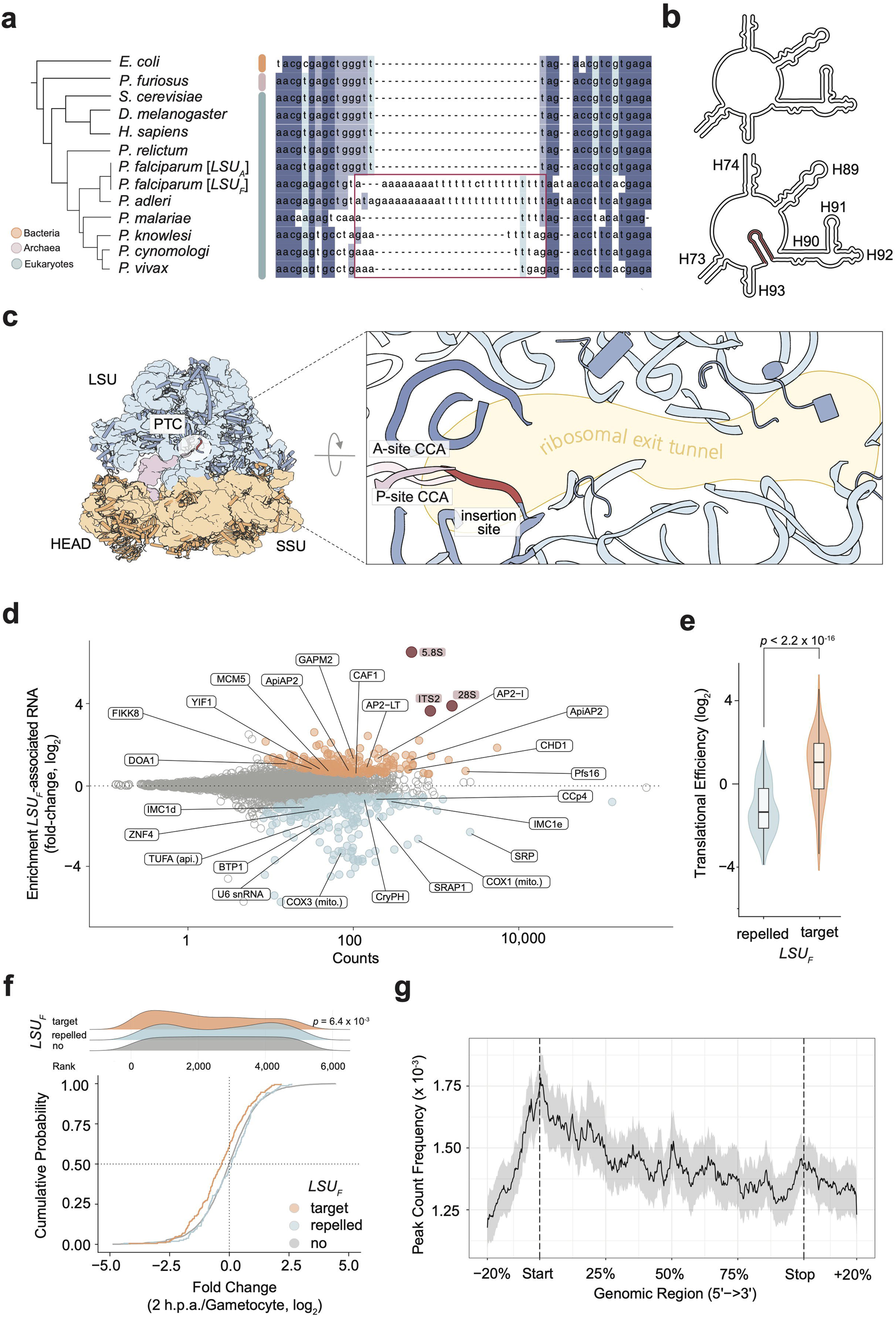
*LSU_F_* engages with mRNA. (a) Multiple sequence alignment of the PTC from archaea, bacteria and eukaryotes. The hairpin sequences of *LSU_F_* homologs is highlighted in red. (b) Secondary structure map of the PTC in *LSU_A_* (top) and *LSU_F_*(bottom) rRNA. The hairpin in the *LSU_F_* rRNA is shown in red. H: Helix. (c) Three-dimensional structure of the canonical *P. falciparum* A-type ribosome (left, PDB: 9J9H) and detailed view of the ribosomal exit tunnel (right). The insertion site of the *LSU_F_*PTC hairpin is highlighted in red (see also panel a). (d) Relative enrichment and depletion of mRNAs associated with *LSU_F_*. Red: *LSU_F_* rRNA. Orange: *LSU_F_* targets (*q* ≤ 0.05. fold-change ≥ 1.5); Blue; *LSU_F_*repelled: (*q* ≤ 0.05. fold-change ≤ -1.5). (e) Translational efficiencies in wild-type gametocytes for *LSU_F_* repelled (*n* = 158) and *LSU_F_*target (*n* = 187) mRNA transcripts. *p*-values were calculated with an unpaired Wilcoxon Signed Rank Test. (f) Empirical cumulative distribution function of changes in translational efficiency for *LSU_F_*target, *LSU_F_* repelled and other transcripts between gametocytes and gametes at 2 h.p.a. The distribution of ranked changes in translational efficiency for the three classes of mRNAs is shown above. (g) Metagene plot showing the average enrichment of *LSU_F_* associated mRNA reads across mRNA transcripts. The grey area indicates 95% confidence interval. Start: start codon. Stop: stop codon.

Within *LSU_F_* rRNA, we identified multiple characteristic long-range interactions predicted from the secondary structure model, including those between the 5.8S and 28S, multiple interactions in domain II, the PTC (Fig. 2e), and between the *LSU_F_* 28S and the canonical *SSU_A_*18S rRNA (Fig. 2h). In addition, we also detect specific interactions within the retained *LSU_F_* ITS2 ‘foot’ structure, yet only minimal interaction of this region with either the *LSU_F_* 5.8S or 28S rRNA (Fig. 2h). Hence, *LSU_F_* ITS2 constitutes an independently folding rRNA domain, in accordance with ribosomal assembly intermediates. Collectively, these findings reveal that *LSU_F_*rRNA has retained and further evolved both conserved and unique characteristics of rRNA composition and structure, thereby possibly contributing to the formation of a nuclear-encoded, highly divergent ribosome core. These include conserved rRNA modifications and overall folding, the evolution of vast expansion segments as well as the loss of an encoded 18S and ITS2 splicing, differences that overall amount to those usually seen only in rRNA of very distantly related organisms (Supplementary Fig. 2e).

### *LSU_F_* engages with mRNA

Modelling of the 2D structure and RIC-seq confirmed that *LSU_F_* retains a structured PTC. In agreement with its catalytic role in protein synthesis, the PTC is the most ancient and part of the rRNA core, universally conserved across all domains of life^41^ (Fig. 3a, Supplementary Fig. 2e). While this also holds true for the canonical *LSU_A_*rRNA of *P. falciparum,* we find that the PTC of *LSU_F_* homologs include a preemptive hairpin structure (24 nt in length in *P. falciparum*) extending from between helix 92 and 93 (Fig. 3b). This PTC hairpin is absent from the large subunit rRNAs encoded at the corresponding *P. relictum* rDNA locus and coincides with the loss of the SSU rRNA and retention of the ITS2 ‘foot’ in *LSU_F_* homologs (Fig. 3a).

Mapping the insertion site on the atomic structure of the canonical *P. falciparum* ribosome shows that this hairpin disrupts the structure of the PTC, directly interferes with the CCA tRNA 3’ ends located at the A- and P-sites, and can entirely block the ribosomal exit tunnel (Fig. 3c, Supplementary Fig. 4a). Sterically inhibiting a nascent peptide chain via this hairpin structure is prohibitive for protein synthesis and raises the question whether *LSU_F_*can interact with mRNA. We therefore used a pooled, antisense-oligo pulldown approach (‘raPOOL’)^48^ targeting the 28S sequence of either the canonical *LSU_A_* or variant *LSU_F_*rRNA to identify cognate mRNA transcripts in wild-type gametes at 2 h.p.a. (Supplementary Fig. 4b, Supplementary Table 4). In parallel, we performed ribosome profiling (Ribo-seq) of wild-type gametes at 2 h.p.a. (Supplementary Fig. 4c,d, Supplementary Table 5). A direct comparison of the two orthogonal approaches shows a positive relationship in read coverage between mRNA fragments that were pulled-down with *LSU_A_* 28S-targeting oligos (Supplementary Fig. 4e), ribosome protected fragments (RPFs, Supplementary Fig. 4f) and translational efficiencies (Supplementary Fig. 4g) identified via Ribo-seq^49^. This confirmed that raPOOL can recapitulate *bona fide* mRNA-ribosome interactions in a LSU rRNA specific manner.

The most highly enriched transcripts in the *LSU_F_*rRNA pulldown include the *LSU_F_* 5.8S rRNA and ITS2, providing further evidence that the latter remains attached in a mature *LSU_F_*(Fig. 3d). In addition, we identify 191 RNA transcripts that are significantly (*q* ≤ 0.05. fold-change ≥ 1.5) enriched in the pull-down fraction (*LSU_F_* ‘targets’), and 215 that are significantly depleted (*LSU_F_* ‘repelled‘, *q* ≤ 0.05. fold-change ≤ -1.5, Fig. 3d, Supplementary Table 4). Similar to mRNA transcripts that are depleted from the *LSU_A_*rRNA pulldown (Supplementary Table 4), *LSU_F_* repelled transcripts include spatially segregated, organellar transcripts encoded by the mitochondrial and apicoplast genomes in addition to different ncRNA and multiple proteins of the inner membrane complex (Fig. 3d). *LSU_F_*targets do not share any specific functional relationship but include several ApiAP2 transcription factors, a FIKK kinase (FIKK8) and the parasitophorous vacuole membrane protein S16 (Pfs16), among others (Supplementary Table 4,6). However, we find that a unifying pattern of *LSU_F_*target mRNA transcripts is their high translation efficiencies (Fig. 3e). Indeed, *LSU_F_* targets feature among the highest translational efficiencies in the gametocyte stage prior to transmission, but not at 2 h.p.a (Fig. 3e, Supplementary Fig. 4g). In accordance, translational efficiencies of *LSU_F_* targets are significantly more often downregulated (*χ*^2^, *p* = 6.4x10^-3^) during host-to-vector transmission (Fig. 3f, Supplementary Fig. 4g). This pattern is directly opposite to *LSU_F_*repelled transcripts; translational efficiencies of *LSU_F_* repelled mRNA transcripts in gametocytes are significantly lower than those of *LSU_F_*target mRNAs (Fig. 3e). Despite being significantly more abundant than *LSU_F_* target transcripts in both developmental stages (Supplementary Fig. 4h), they are also among the lowest translated transcripts in both stages (Supplementary Fig. 4g).

Hence, it appears that high rates of translation in the gametocyte stage could be one determining factor that leads to more frequent engagement of these mRNAs with *LSU_F_*, whose synthesis is initiated immediately after parasite transmission.

Read coverage distribution patterns of ribosome-associated mRNA fragments can be used to identify transcript positions and characteristics that lead to biased ribosome localization or stalling^50^. Using the distribution of RNA fragments pulled down in the *LSU_F_* raPOOL experiment as an indicator of *LSU_F_* localization along a mRNA transcript, we find a strong enrichment towards the 5’ end, with the main peak being located at the start codon (Fig. 3g). In contrast, fragments that were pulled down in the *LSU_A_* targeted raPOOL are equally distributed along mRNA coding sequences, a pattern similar to that of RPFs (Supplementary Fig. 4i,j). Taken together, these observations suggest that *LSU_F_* is capable of associating with mRNA, potentially through interactions with the pre-initiation complex at the start codon. However, *LSU_F_* appears to be arrested at the start codon and unable to recruit the tRNAs necessary for efficient translation elongation, consistent with the presence of a disruptive A-site hairpin. Importantly, we identified *LSU_F_* targets at 2 h.p.a., at a time when translation efficiencies of their target mRNAs are already repressed and many mRNAs feature higher translation efficiencies than *LSU_F_* targets. Hence, while high translation efficiency may serve as the initial determinant of *LSU_F_* binding, these findings also suggest that *LSU_F_*remains stably associated at the start codon of its cognate mRNA once bound, possibly leading to the repression of protein synthesis.

### *LSU_F_* represses mRNA translation

To provide functional evidence for a possibly repressive role of *LSU_F_*, we generated a *LSU_F_* knock-out cell line using CRISRP/Cas9 genome editing (‘Δ*LSU_F_*’, Supplementary Fig. 5a,b). Δ*LSU_F_*parasites develop from asexually replicating parasites into gametocytes following a similar phenotypic and transcriptional trajectory as wild-type parasites (Supplementary Fig. 5c-f, Supplementary Table 7). Similarly, a direct comparison of the WT and Δ*LSU_F_*transcriptomes identified only minimal changes at 2 h.p.a. (Supplementary Fig. 6a,b, Supplementary Table 8). Hence, *LSU_F_*rRNA appears to be dispensable for the re-initiation of development, allowing for an unbiased functional characterization of *LSU_F_* during transmission.

At the translational level, ribosome profiling of Δ*LSU_F_* parasites at 2 h.p.a (Supplementary Fig. 6c) shows a similar transition as wild-type parasites that is distinct from the translational program observed in mature gametocytes (Supplementary Fig. 6d, Supplementary Table 8). In contrast to the transcriptional changes, a direct comparison of translational identified 788 transcripts with significantly (*q* ≤ 0.05) higher (‘Δ*LSU_F_*^UP^*’*) translational efficiencies, whereas 784 were significantly lower (‘Δ*LSU_F_*^DOWN^*’*) (Fig. 4a, Supplementary Fig. 6e, Supplementary Table 9). The encoded products of neither Δ*LSU_F_*^UP^, nor Δ*LSU_F_*^DOWN^ mRNAs are significantly enriched in a functional category (Supplementary Table 6). However, Δ*LSU_F_*^UP^ and Δ*LSU_F_*^DOWN^ mRNAs follow two distinct translational programs during transmission when in a wild-type context; Δ*LSU_F_*^UP^ feature high translational efficiencies prior to transmission (Fig. 4b), and their translation is repressed after (Fig. 4b, Supplementary Fig. 6f,g Supplementary Table 9). Among those mRNAs are transcripts important for the gametocyte stage, including DOZI and CITH as well as Pfs16 and FIKK8, which we identified also among the *LSU_F_* target transcripts (Fig. 3d, 4a). Conversely, Δ*LSU_F_*^DOWN^ mRNA exhibit low translation rates that are significantly lower than those of Δ*LSU_F_*^UP^ prior to transmission (Fig. 4b,c), but their translation is de-repressed once development is re-initiated (Fig. 4b, Supplementary Fig. 6g,g Supplementary Table 9) and include mRNAs that were also among the *LSU_F_* repelled transcripts, including components of the inner membrane complex (Fig. 3d, 4a). However, following *LSU_F_* deletion, the translation of Δ*LSU_F_*^UP^ mRNAs is not repressed, whereas translation of Δ*LSU_F_* ^DOWN^ mRNAs fail to be initiated after transmission (Fig. 4c). Thus, in a direct comparison at 2 h.p.a., the translational efficiencies of Δ*LSU_F_*^UP^ and Δ*LSU_F_*^DOWN^ appear to be relatively up- or downregulated, respectively, suggesting that the natural switch of the translational program occurring during transmission is partially disrupted in Δ*LSU_F_* parasites (Fig. 4a,c).

**Figure 4.**
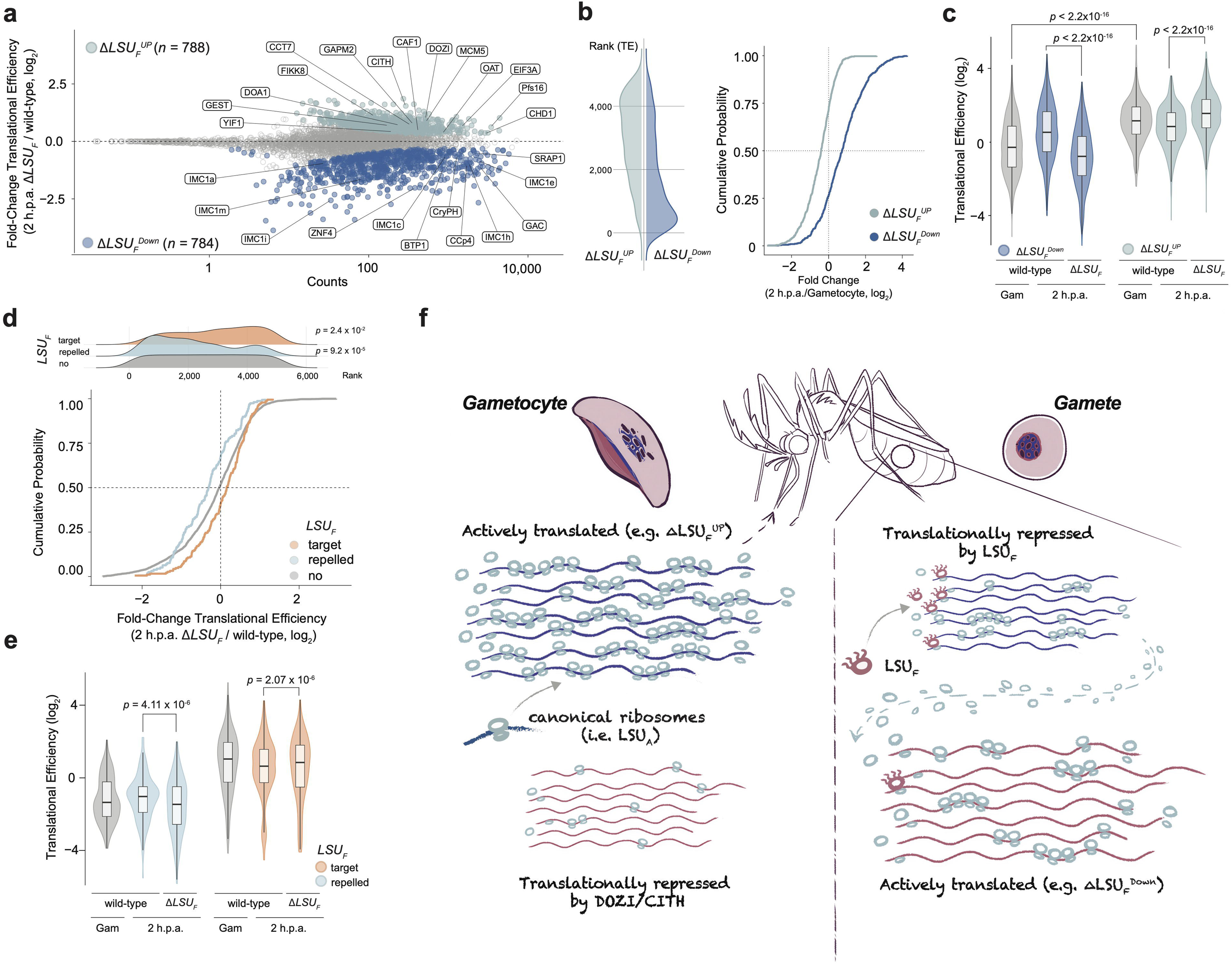
*LSU_F_* represses mRNA translation. (a) Changes in translational efficiencies between wild-type and Δ*LSU_F_* parasites at 2 h.p.a. detected by ribosome profiling. mRNA transcripts with significantly higher (Δ*LSU_F_*^UP^) or lower (Δ*LSU_F_*^DOWN^) translational efficiencies are shown in blue and red, respectively. (b) Left: Transcriptome-wide ranking of translational efficiencies in gametocytes for mRNA transcripts with significantly increased (Δ*LSU_F_*^UP^, top) or decreased (Δ*LSU_F_*^DOWN^, bottom) rates of translation at 2 h.p.a. following *LSU_F_*deletion. Right: Cumulative probability of changes in translational efficiency for Δ*LSU_F_*^UP^ and Δ*LSU_F_*^DOWN^ mRNA transcripts in wild-type parasites between gametocytes and gametes at 2 h.p.a. (c) Translational efficiencies in wild-type gametocytes, wild-type gametes (2 h.p.a.) and Δ*LSU_F_* gametes (2 h.p.a.) for Δ*LSU_F_*^UP^ (left) and Δ*LSU_F_*^DOWN^ (right) mRNA transcripts. *p*-values are calculated with a paired Wilcoxon Signed Rank Test. Δ*LSU_F_*^UP^: *n* = 788; Δ*LSU_F_*^DOWN^: *n* = 784. (d) Empirical cumulative distribution function of changes in translational efficiency for *LSU_F_*target, *LSU_F_* repelled and other transcripts between wild-type and Δ*LSU_F_* gametes at 2 h.p.a. The distribution of ranked changes in translational efficiency for the three classes of mRNAs is shown above. (e) Translational efficiencies in wild-type gametocytes, wild-type gametes (2 h.p.a.) and Δ*LSU_F_* gametes (2 h.p.a.) for *LSU_F_* repelled (left) and *LSU_F_*target mRNA transcripts (right). *p*-values were calculated with a paired Wilcoxon Signed Rank Test. *LSU_F_* repelled: n = 158; *LSU_F_*target: *n* = 187. (f) Proposed model illustrating how *LSU_F_*facilitates the transition of the translational program during *P. falciparum* transmission. Through its preferential binding to and repression of mRNAs that were already highly translated prior to transmission, canonical ribosomes can more frequently engage with newly de-repressed mRNAs that ensure continuation of mosquito stage development.

Integration of the *LSU_F_* raPOOL data shows that *LSU_F_* target mRNAs are significantly enriched (*χ^2^*, *p* = 2.4x10^-2^) among mRNAs whose translation efficiencies increase in Δ*LSU_F_* at 2 h.p.a. *LSU_F_*repelled mRNAs on the other hand are significantly overrepresented *χ^2^*, *p* = 9.2x10^-5^) among transcripts with decreased translation rates (Fig. 4d, Supplementary Fig. 6h). It accordance, we find a similar pattern of translational dynamics for *LSU_F_* targets as we see for Δ*LSU_F_*^UP^, i.e. translational repression in wild-type parasites after transmission, but a lack thereof in Δ*LSU_F_* cells (Fig. 4e). In contrast, translation of *LSU_F_* repelled mRNAs increases during transmission in wild-type cells, but fails to do so once the *LSU_F_*rRNA is disrupted, similar to what is observed for Δ*LSU_F_*^DOWN^ transcripts (Fig. 4e). This implies that the translational repression of Δ*LSU_F_*^UP^ is a direct function of *LSU_F_* binding, and that a lack thereof impedes the initiation of the early mosquito stage translational program, including the translational activation of Δ*LSU_F_*^DOWN^.

Despite their large differences in translational efficiencies, Δ*LSU_F_*^UP^ and Δ*LSU_F_*^DOWN^ mRNAs feature similar levels of global transcript abundance in wild-type mature gametocytes (Supplementary Fig. 6i). Because the canonical *LSU_A_* rRNA is not substantially upregulated during transmission of wild-type parasites (Supplementary Fig. 1e,f), the release of mRNAs from translational repression alone could therefore explain the observed translational dynamics of relative up- and downregulation (Fig. 4c) due to more mRNAs being permissible to translation and a relative dilution of the available canonical ribosome pool. However, we find that even though Δ*LSU_F_* parasites can re-initiate development (Supplementary Fig. 6b,d), Δ*LSU_F_*^DOWN^ mRNAs do not reach the same level of translational efficiencies as in wild-type gametes (Fig. 4a,c). This observation suggests that previously translationally repressed mRNAs are unable to effectively ‘compete’ for what is likely a limiting pool of ribosomes, and that *LSU_F_* constitutes an additional regulatory layer governing the reprogramming of translation after transmission (Fig. 4f). In order to act on a broad range of mRNAs without the necessity of additional specificity factors, the most parsimonious model of how *LSU_F_*can facilitate such a global redistributing of available resources is by binding to the TSS of mRNAs that were already highly translated prior to transmission (i.e. Δ*LSU_F_*^UP^), which in return leads to an increased pool of available ribosomes that initiate with newly de-repressed mRNAs (i.e. Δ*LSU_F_*^DOWN^, Fig. 4f). Here, only few numbers of repressive *LSU_F_* ribosomes might be necessary to impede the linear processing of polysomes along a translated mRNA, akin to how translation rates can decrease due to ribosome stalling^51^. Although the *LSU_F_*-*SSU_A_* interactions identified via RIC-seq and the enrichment at the start codon provide some evidence that *LSU_F_* could bind mRNA via the pre-initiation complex, the exact nature and structure of how *LSU_F_* interacts with the broader canonical translational machinery remains to be determined. Importantly though, the finding that *LSU_F_* appears to remain attached once engaged with a target mRNA suggests that the overall low numbers of *LSU_F_* (compared to the canonical *LSU_A_*) could be rapidly depleted, further aiding the activation of de-repressed mRNAs. In addition, given that both DOZI and CITH are among Δ*LSU_F_*^UP^ mRNAs, *LSU_F_* could also directly act to repress the repressor, potentiating the effect of the translational transition during transmission.

A functionally unspecific, global switch of protein synthesis where translational efficiency alone can function as the determining factor for *LSU_F_* engagement might be especially efficient in this system where 1) the discrepancies of translation rates between two regulated mRNA pools requiring opposing translation regulation are large, and 2) where the transitions between developmental stages is not intrinsically regulated, but triggered by sudden environmental changes. Most importantly though, given the absence of transcriptional regulation or targeted mRNA degradation (e.g. of Δ*LSU_F_*^UP^, Supplementary Fig. 6a), rapidly redistributing already available resources solely through *LSU_F_* transcription might be more cost effective than engaging in *de novo* biosynthesis of canonical ribosomes to accommodate the translation of newly de-repressed mRNAs.

## DISCUSSION

Our study identifies a genomically-encoded, variant rRNA allele that provides an inherent regulatory capacity for a functionally divergent, repressive ribosome in the human malaria parasite. For *P. falciparum*, this facilitates the attenuation of the translational program of one developmental stage and allows for the transition into a new one that supports the continuation of development. While we focused on the experimentally accessible human-to-mosquito transmission stage, the additional periods of *LSU_F_*transcription after ookinete-to-oocyst transition and following mosquito-to-host transmission (during development in the human liver) suggest that *LSU_F_* could facilitate a similar switch of translational programs during major developmental transitions across the parasite lifecycle. However, the essentiality and exact function at these developmental stages remains to be elucidated.

Compositional ribosome heterogeneity around the same rRNA has been widely observed across organisms of all kingdoms of life^17^, providing the translational machinery with a means to adapt to different cellular states^52^. Our data show that functional divergence of different ribosome populations can emerge from the incorporation of a distinct (non-)catalytic rRNA core itself. We show that this extends the regulatory ability of ribosome populations on two temporal scales. On the short-term, the precisely controlled co-transcription of variant rRNA alleles allows them to be selectively employed to differentially alter the translational output of subsets of mRNA, e.g. in the context of a developmental transition. On the long-term, our data provide evidence that separate rDNA loci can follow independent evolutionary trajectories to evolve a new function for global translational regulation, analogous to how novelty can arise for protein coding genes after a gene duplication. In prokaryotes, the transcription of distinct rRNA operons has been linked to the translational response to stress^13,14^, and variant rRNA expression has also been associated with different pathologies in humans^8,9^. However, we find that instead of providing a ‘specialized’ function^17,53^ to its encoded LSU that enhances or maintains protein synthesis under changing environmental or cellular states^54^, the *LSU_F_*rRNA evolved an entirely new and opposite, repressive role. Thus, *LSU_F_*comprises an additional class of a rRNA-derived macromolecular complex with a repressive role in mRNA translation. Mechanistically, it is plausible that this repressive function is encoded in the highly divergent structure of the PTC. This unique modification is likely able to entirely disrupt peptide chain elongation in the exit tunnel. Yet whether it is the first structural feature providing the repressive function is unknown. Accordingly, if the emergence of any additional diverging features in *LSU_F_* is a result of selective pressure to enhance its repressive function or emerged from genetic drift due to the absence of selective pressure on *LSU_F_* to maintain its protein synthesis capability remains to be addressed. More importantly however, because *LSU_F_* acts simultaneously and antagonistic to canonical ribosomes within the same cell, this data establishes that *LSU_F_* serves as a dominant-negative regulator and exemplifies that ribosomes can diverge from their universally conserved function in mRNA translation.

Taken together, we identify a repressive ribosome population whose regulatory function in protein synthesis is encoded by an independently evolved rRNA allele, providing the conceptual foundation of variant rRNA-mediated ribosome heterogeneity as an additional, inherent layer of gene regulation across biological systems.

## ACKNOWLEDGEMENTS

We thank Sylvain Gandon for sharing the *P. relictum* genome assembly. Research in the laboratory of S.B. is supported by an ERC Starting Grant (# 947819) and baseline funding of the Institut Pasteur and INSERM. J.M.B. is supported by the French National Research Agency ANR-21-CE15-0002-02 (ApiMORCing) and ANR-21-CE15-0010-01 (PlasmoVarOrg). V.T. acknowledges support from Wellcome Trust (# 325247/Z/25/Z). The work in the laboratory of S.S. is funded by an ERC Consolidator Grant (# 101000970). S.B., S.S. and M.S.B. were supported by the Pasteur-Weizmann Foundation.

## METHODS

### Parasite cell culture

Asexual blood-stage *P. falciparum* parasites were cultured in human RBCs (obtained from the Etablissement Francais du Sang with approval number HS 2021-24819) in RPMI-1640 medium (Thermo Fisher no. 53400-025) supplemented with 10% v/v Albumax I (Thermo Fisher no. 11020039), hypoxanthine (0.1□mM final concentration, CC-Pro no. Z-41-M) and 10□mg gentamicin (Sigma no. G1397-10ML) at 4% haematocrit and under 5% O_2_, 3% CO_2_ at 37°C. Parasite development was monitored by microscopic observation of Giemsa-stained thin blood smears. For synchronization, late-stage parasites were enriched by plasmion flotation followed by ring-stage enrichment via sorbitol (5%) lysis 6 hours later, resulting in an average parasite population age of 3 h.p.i (± 3 hours).

### Generation of Gametocytes

Gametocytes were obtained following the protocol by Fivelman *et al.*^64^. Briefly, synchronous asexual, late-stage parasites (30-35 h.p.i.) were concentrated to ∼2.5% parasitemia and 2.5% hematocrit using plasmion flotation. The next day, 75% of the spent culture medium was replaced with fresh medium, and the ring-stage parasites (∼10-15% parasitemia) were left to develop into trophozoites at high parasitemia for an additional 24h. The culture was then diluted in fresh media to 3% parasitemia and kept for an additional 24 hours. The growth medium of the resulting high-parasitemia, ring-stage culture was then replaced with RPMI supplemented with 5% human serum, 5% Albumax, 0.1 mM hypoxanthin, 10 mg Gentamicin and 50 mM N-acetylglucosamine (NAG, Sigma no. A3286). Growth media was then changed daily for 5 days with the addition of NAG, and then without NAG for an additional six days. For sample collection, all gametocytes were first enriched using NycoPrep gradient. In brief, a 5mL Nycodenz cushion (5 mM Tricine [Sigma Aldrich no. T5816], 14.1% w/v Nycodenz [Serumwerk no. 18003], 75 mM NaCl [Sigma Aldrich no. S9888], pH 7.2) was prepared in a 15mL falcon. Next, 1mL packed RBCs were resuspended in 10mL of suspended animation (SA) buffer (RPMI1640 [Gibco no. 11835-063], 25mM HEPES [Gibco no. 15630-056], 5% fetal calf serum [Gibco no. A5209401] and 4 mM sodium bicarbonate [Sigma Aldrich no. S5761], pH 7.2), pipetted on top of the Nycodenz cushion without disturbing the surface and centrifuged (20 min, 800 x g, 37°C and no break). The grey interphase formed by gametocytes was retained and directly mixed in 8 mL SA buffer, and centrifuged (5 min, 975 x g, 37°C). The supernatant was removed and the purified gametocyte pellet used for downstream sampling or activation.

### Generation of gametes (Gametocyte activation)

*P. falciparum* gametocytes were activated to form gametes following the ookinete culture protocol described in ref.^32^, with some minor modifications. Briefly, mature, stage V gametocytes were separated from uninfected RBC at day 11 using NycoPrep density centrifugation maintaining all reagents and solutions at 37°C to prevent premature activation (see above). For activation, Gametocytes were spun down (37°C and 800 x g) in 50 ml tubes, followed by removal of growth medium. A volume of room temperature (∼23°C, RT) human serum equal to the volume of the infected RBC pellet was added (i.e. defining the time point of gametocyte activation, 0 h.p.a), cells were resuspended by gentle flicking and incubated for 30 min at RT. Next, 5X RBC pellet volumes of room temperature ookinete medium (RPMI-1640, 25 mM HEPES, 2 mM L-glutamine, 2 g/L sodium bicarbonate, 50 mg/L hypoxanthine, pH 8.2)^65^ were added, mixed, and incubated for an additional 90 min at RT, resulting in the formation of 2 h.p.a. gametes.

### Generation of cell lines

The Δ*LSU_F_* cell line was generated via CRISPR/Cas9 induced double-strand break (DSB) in the P. falciparum wild-type strain NF54. Two plasmids were co-transfected to remove 1,044 bp of the *LSU_F_* 28S rRNA (PF3D7_0801100: 4,389 – 5,433), encompassing a region within domain V that encodes the PTC as well as the L1 stalk) and insert a selectable resistance cassette encoding a human dihydrofolate reductase (hDHFR) via homologous recombination (Supplementary Fig. 5a).

The first plasmid (pDC2-Cas9-yDHODH-*LSU_F_*), encodes the SpCas9 nuclease, a selectable marker (yeast dihydroorotate dehydrogenase, yDHODH) and a single guide RNA (sgRNA) expression cassette^30,66,67^ (Supplementary Fig. 5a). A sgRNA target sequence containing a NGG target sequence located inside the 1,044 bp that were targeted for excision was designed using CHOPCHOP^68^. pDC2-Cas9-yDHODH was digested with BbsI and the *LSU_F_* rDNA specific sgRNA (sgRNA-*LSU_F_*) was cloned into the digested backbone, yielding pDC2-Cas9-yDHODH-*LSU_F_*.

The second plasmid (pUC57-LSU_F_-hDHFR), contains the homologous repair template to insert the selectable hDHFR resistance cassette into the *LSU_F_* rDNA locus. First, two 400 bp-long DNA homology regions separated by AvrII and NcoI restriction sites were synthesized by GenScript and cloned into a pUC57 plasmid, yielding pUC57_*LSU_F_*-HR. The homology regions are identical to the upstream (PF3D7_0801100: 3,989 – 4,389) and downstream (PF3D7_0801100: 5,433 – 5,833) sequences flanking *LSU_F_*rDNA sequence targeted for excision. Next, the hDHFR resistance cassette encoding a 5’ calmodulin (PF3D7_1434200) promoter, the hDHFR coding sequence and a histidine-rich protein 2 (PF3D7_0831800) 3’UTR was amplified by PCR from pL7-eGFP^69^ using primers hDHFR_*LSU_F_*_cloning_F and hDHFR_*LSU_F_*_cloning_R. pUC57_*LSU_F_*-HR was digested with AvrII and NcoI (i.e. separating the upstream and downstream homology regions) and the hDHFR PCR product was cloned in, yielding pUC57-*LSU_F_*-hDHFR (Supplementary Fig. 5a).

Cloning and plasmid amplification were performed using the In-Fusion HD cloning kit (Clontech # 639649) and XL10-Gold ultracompetent *Escherichia coli* (Agilent Technologies # 200315) following the manufacturer’s protocol. All plasmids were verified by Sanger sequencing. Prior to transfection, pUC57-LSUF-hDHFR was linearized by ScaI digestion, purified via phenol-chloroform extraction, and co-precipitated in ethanol together with pDC2-Cas9-yDHODH in a 3:1 molar ratio. Approximately 90 µg of linearized pUC57-*LSU_F_*-hDHFR and 30 µg of pDC2-Cas9-yDHODH were used for transfection. The two plasmids were co-transfected into wild-type ring-stage *P. falciparum* (strain NF54) parasites by electroporation following previously published protocols^69^. To select for pDC2-Cas9-yDHODH uptake, transfected parasites were cultured with 1.5□μM DSM1 (BEI Resources # MRA-1161). After drug-resistant parasites emerged approximately 4 weeks post-transfection, integration of the hDHFR selection marker into the F-type rDNA locus was positively selected through the addition of 5 nM WR99210 (Jacobus Pharmaceuticals). Integration of the resistance cassette was first verified using primers *LSU_F_*_upstream_F and *LSU_F_*_downstream_R. Individual clones of the transfected parasite population were then obtained via serial dilution and initially screened via PCR amplification and Sanger sequencing of the targeted genomic locus using primers *LSU_F_*_upstream_F and *LSU_F_*_downstream_R. Positive clones were further verified by full genome sequencing on the MinION (Oxford Nanopore Technologies) as described previously^12^. In brief, asexual parasites were collected by saponin lysis (0.075% saponin in DPBS) and the cell pellet was washed twice with ice-cold DPBS, snap-frozen and stored at -80°C. gDNA was extracted using the Qiagen DNeasy Blood & Tissue Kit (Qiagen # 69504) and purified by phenol/chloroform extraction and ethanol precipitation. Libraries were prepared using the SQK-RAD004 kit and sequenced on a FLO-MIN106 flow cell on a MinION sequencer. Basecalling was performed using the fast model in Guppy (v 6.5.7) implemented in MinKNOW (v23.04.5). The expected hDHFR resistance cassette was manually integrated at the *LSU_F_* rDNA locus of the reference *P. falciparum* (PlasmoDB v64^70^) genome and annotation. Raw fastq reads were mapped to the modified genome containing the hDHFR cassette in the *LSU_F_* rDNA locus using minimap2^71^ with option ‘-x map-ont’. The resulting alignments were visualized using the Integrative Genomics Viewer^72^ (Supplementary Fig. 5b). Of note, given the proximity of the *LSU_F_* rDNA locus to the left end of chromosome 8, only one clone (out of 20) correctly integrated the recombination template and maintained the downstream chromosome region. The remaining clones integrated the recombination template, followed by *de novo* addition of telomere repeats, thus dropping the remaining ∼90 kb (encompassing nine protein coding genes, two pseudogenes and one H/ACA snoRNA) located on the left arm of chromosome 8

### Total RNA extraction

Parasite cultures were spun down (at 37°C in the case of gametocyte cultures, at room temperature in the case of asexual or post-activation gamete cultures) at 800 x g, followed by removal of growth medium. For asexual parasites, RBCs were lysed by resuspending the infected blood pellet in 10 volumes of 0.075% saponin (Sigma no. S790) in Dulbecco’s phosphate-buffered saline (DPBS, Thermo Fisher no. 14190-144) at 37°C. Parasites were pelleted by centrifugation (2,000 x g, 4 min) and immediately resuspended in TRIzol (Thermo Fisher no. 15596026) after removal of the RBC lysate supernatant.

For gametocytes, uninfected RBC were separated using NycoPrep (see above), and the parasite pellet was directly resuspended in 700 μL TRIzol without removal of the RBC via saponin lysis. For gametes, the cell pellet was also directly resuspended in 700 μL TRIzol. Total RNA was obtained by chloroform extraction followed by co-precipitation with GlycoBlue (Thermo Fisher no. AM9516) in isopropanol. Precipitated RNA was washed with 75% ethanol and reconstituted in nuclease-free water.

### qPCR

Total RNA was extracted from parasites as described above. Contaminating genomic DNA was removed using the TURBO DNA-free Kit (Thermo Fisher #AM1907) following the manufacturer’s instructions, and reverse transcription on the entire transcriptome was performed using the ProtoScript II First Strand cDNA Synthesis Kit (NEB # E6560L) with the provided Randomized Primer Mix. *LSU_F_* and *LSU_A_* rRNAs were amplified in technical triplicates using Power Sybr Green PCR Master Mix (Thermo Fisher # 4367659) and primers LSU-F_F/LSU-F_R and LSU-A_F/LSU-A_R on a BioRad CFX qPCR machine. All experiments included a no-RT control and a no-template control. Serial dilutions of genomic DNA were amplified alongside the experimental samples to generate a standard curve, from which mean starting quantity (SQ-mean) values for the technical replicates were derived.

### mRNA sample preparation and sequencing

mRNA was enriched from total RNA using the Dynabeads mRNA Purification Kit (Thermo Fisher no. 61006). Sequencing libraries were prepared with the NEBNext Ultra II Directional RNA Library Prep Kit (NEB no. E7760S) according to the manufacturer’s instructions for fragment sizes of 300 bp. Libraries purity and size distributions were assessed on a TapeStation 4150 (Agilent Technologies) and quantified by qPCR usint the KAPA Library Quantification Kit (Roche no. KK4844). Final libraries were sequenced with a 2x150 bp paired-end layout on an Illumina NextSeq 2000 platform. All total RNA and mRNA sequencing experiments were performed with three independent biological replicates.

### mRNA sequencing analysis

Sequencing reads were demultiplexed with bcl2fastq2 (version 2.20, Illumina), followed by the removal of sequencing adaptors and low-quality reads using trimmomatic^73^. Reads were then aligned to the *P. falciparum* genome (version 64)^23,70^ using STAR^74^ with default settings and only retaining uniquely mapping reads (option --outFilterMultimapNmax 1). PCR duplicates were removed using samtools’ ‘fixmate’ and ‘markdup’ ^75^, and correctly paired reads were filtered using samtools ‘view’ (option -f 0x2). Gene counts were calculated using htseq-count^76^ using alignments that fall within exons (option ‘-t exon’) and differentially expressed genes and log_2_ fold-changes were identified in R^77^ using DESeq2^78^. Alignments were visualized using the Integrative Genomics Viewer^72^.

The approximation of the developmental age by comparison to scRNA-seq (Supplementary Fig. 5f) was calculated as described previously^12^. In brief, for each gene, the average FPKM across all three replicates was calculated and the expression of all genes was correlated to each individual transcriptome of Dogga et al.^63^ in R using cor(method = “pearson”). Resulting Pearson *R^2^* values were plotted over the UMAP1 and UMAP3 coordinates of each individual cell in R using ggplot2^79^.

For the comparison of *LSU_F_* ortholog expression in other *Plasmodium* species (Supplementary Fig. 1c), the raw data were downloaded from SRA using fastq-dump from sra-toolkit^80^. Accession numbers of all samples are listed in Supplementary Table 12. Raw reads were pre-processed, mapped and alignments were filtered as described above. For read mapping and feature counting, the respective reference genome and annotation of each species was downloaded from PlasmoDB (version 67)^70^

### Ribosome profiling

Samples from the same three independent biological replicates of wild-type (strain NF54) and Δ*LSU_F_ P. falciparum* gametes (2 h.p.a.) that were used for mRNA sequencing (see above) were also collected in parallel for ribosome profiling. Gametes were obtained as described above via NycoPrep purification, washed once in ice-cold DPBS, snap-frozen in a dry ice-ethanol bath, and stored at -80°C until processing.

Actively translating ribosomes were purified using the RiboLace Pro kit (Immagina Biotechnology no. RL00P-12) following the manufacturer’s instructions. Shortly, cell pellets were lysed in the provided cycloheximide-containing lysis buffer, supplemented with sodium deoxycholate (1%), RNase-free DNase I (NEB no. M0303S) (5U/mL), and RiboLock Rnase Inhibitor (Thermo Fisher no. EO0381) (200 U/mL), followed by mRNA digestion with the provided nuclease to generate ribosome protected fragments (RPFs). Ribosomes were captured by incubating the digested lysate with functionalized magnetic beads for 70 min at 4°C under slow rotation, and washed twice on ice with the provided wash buffer. RPFs and other remaining RNAs were released by treating the beads with the provided proteinase K solution in wash buffer supplemented with 1% SDS for 75 min at 37°C. The reaction was stopped by mixing with 3X volumes of TRIzol LS and RNA was purified via TRIzol-chloroform extraction as described above. RPFs were separated from other RNAs present in the samples via size-selection on a denaturing polyacrylamide gel. Briefly, RNA obtained through RiboLace was loaded on a 10-well Novex 15% TBE-Urea Gel (Thermo Fisher no. EC6885BOX) alongside a 25-35 bp marker (included in the RiboLace Pro kit) and an ultra-low range molecular weight marker (Thermo Fisher no. 10597012), and ran at 180V for 1.5 hours. The gel was then stained with SYBR Gold (Thermo Fisher no. S11494) in TBE buffer for 5 min, and visualized on a blue-light transilluminator. The gel sections containing RPFs (between approximately 25 and 35 nucleotides in length) were cut with a clean scalpel, finely crushed, and mixed with 300 μL of gel extraction buffer (300 mM sodium acetate, 1 mM EDTA, 0.25% v/v SDS). The buffer-gel suspensions were snap-frozen in dry ice-ethanol bath and kept at -80°C for 30 min, and the RNA was then eluted from the gel by incubating overnight at 25°C under gentle shaking (500 r.p.m.).

The next day, the eluate was separated from the crushed gel through spin-filtration on Costar Spin-X tube filters (Corning no. 8161) and the RNA was co-precipitated with GlycoBlue (Thermo Fisher no. AM9516) by adding 3 volumes of 95% ethanol and incubating at -80°C for at least 30 min. After spinning down (12,000 x *g* for 15 min), RNA pellets were washed with 80% ethanol, air-dryed, and reconstituted in 10 μL of nuclease-free water.

RPFs were end-repaired by incubating with 10 units of T4 PNK (NEB no. M0201S) in 18 μL of 1X T4 PNK buffer in the presence of 20 units of murine RNase inhibitor (NEB no. M0314S), for 30 min at 37°C. 2 μL of 10 mM ATP was then added before extending the incubation by another 30 min at 37°C.

The reaction was stopped by precipitating the RNA with 36 μL of RNAClean XP beads (Beckman Coulter no. A63987) and 60 μL of isopropanol on ice for 20 min. Leftover ATP was removed by washing the beads twice with 80% ethanol, and RPFs were finally eluted in 6 μL of nuclease-free water. Sequencing libraries were prepared from the RPFs with the NEBNext Small RNA Library Prep Set for Illumina (NEB no. E7330S), and further size-selected via Pippin Prep (Saga Science) with a fragment size range of 125-175 bp before sequencing (100 bp single-end) on an Illumina NextSeq 2000 platform.

### Analysis of translation efficiency

Raw reads of ribosome protected fragments were demultiplexed using bcl2fastq2 (version 2.20, Illumina) followed by the removal of sequencing adapters and low-quality reads using trimmomatic^73^. The reads were then aligned to the *P. falciparum* reference genome (version 64)^23,70^ using STAR^74,75^ with default settings and only retaining uniquely mapped reads. PCR duplicates were removed using samtoos ‘rmdup’. For the calculation of steady state mRNA levels that serve as the basis for the determination of translation efficiency, gene counts from the mRNA sequencing were calculated as described above (see mRNA sequencing and analysis) using htseq-count, but only considering alignments falling within protein coding sequences (option ‘-t CDS’). Gene counts of ribosome protected fragments were calculated in an identical manner, i.e. using htseq-count and only considering alignments falling within protein coding sequences (option ‘-t CDS’). FPKM values (fragments per kilobase of exon per one million mapped reads) for mRNA and RPF gene counts were calculated using DESeq2, using the total length of the coding sequence of each gene as the length normalization factor. Translation efficiencies were then calculated as TE = FPKM_RPF_/FPKM_mRNA_. Differential RPF occupancy was calculated as described for differentially expressed genes in DESeq2 (see above). Differential translation efficiencies were calculated using the raw gene counts of RPF and mRNA in R^77^ using deltaTE^81^. Translational efficiencies of wild-type *P. falciparum* gameotcytes were taken from ref. ^30^

### Generation of 2D rRNA structures and expansion nomeclature

Sequences from *P. falciparum LSU_F_* and *LSU_A_* rRNA as well as outgroup *Plasmodium* species (*P. relictum, P. praefalciparum, P. malariae, P. vivax, P. knowlesi, P. cynomologi, P. billcollinsi, P. blacklocki, P. adleri* and *P. gaboni*) were used to generate a master alignment using muscle5^39^. Due to prominent low complexity regions in the sequences, a total of 100 diversified sequence alignments were generated as “muscle5 -diversified -replicates 100” and then averaged using “muscle5 -maxcc”. The resulting initial alignment was then manually curated to remove large insertions and iteratively realigned using the same approach to produce a high confidence consistent core alignment, corresponding well with the *LSU_A_*core. The full *LSU_F_* sequence was then realigned against this *LSU_A_* cropped core, once again using diversified muscle, to produce the *LSU_F_* to *LSU_A_* alignment. This alignment was used as a reference for 2D structure generation using xrna 1.2.0 with secondary structures of expansion regions predicted using the ViennaRNA 2.6.3 software suite^40^. All identified expansion segments were labeled as either a further expansion of existing segments following the mammalian nomenclature (ex. ES7A-1), or as an *LSU_F_*specific expansion of underlying bacterial helices (ex. H78-ES1).

### RNA *in situ* conformation sequencing (RIC-seq) and analysis

RIC-seq experiments were performed for three independent biological replicates. *P. falciparum* (strain A11) gametes at 2 h.p.a. were obtained as described above via NycoPrep purification and parasites were washed once with DPBS at room temperature. Cells were cross-linked by adding methanol-free formaldehyde (Thermo Fisher no. 28908, final concentration of 1%) and incubated for 10 min under slow rotation at RT. The cross-linking reaction was quenched by adding glycine to a final concentration of 125 mM and cells were incubated for an additional 5 min. Cross-linked parasites were centrifuged (3,250 x *g*, 5 min) washed once with DPBS, snap-frozen in a dry ice-ethanol bath, and stored at -80°C until processing for RIC-seq. RIC-seq was performed as described in Cao et al.^46^ with minor modifications to adapt it to *P. falciparum* ribosomal RNA. Briefly, a preliminary titration of the MNase concentration was performed on test samples to identify the optimal digestion conditions for RIC-seq (Supplementary Fig. 3b), and 6x10^-6^ units of MNase per μg of total RNA present in the sample was chosen as the ideal amount. RIC-seq was performed as described by the original authors, but omitting the rRNA depletion step prior to the streptavidin-based pulldown of pCp-labelled RNA fragments. After eluting pCp-labelled fragments from the beads, 3 volumes of TRIzol LS (Thermo Fisher no. 10296010) were added to the eluate and RNA was purified via TRIzol-chloroform extraction as described above. Sequencing libraries were prepared with the NEBNext Ultra II Directional RNA Library Prep Kit (NEB no. E7760S) following the protocol for use with rRNA Depleted FFPE RNA and sequenced (2x150 bp paired-end) on an Illumina NextSeq 2000 platform.

For the RIC-seq analysis, we followed the ‘RICpipe’ workflow described in ref.^46^ (https://github.com/caochch/RICpipe), with multiple adjustments to increase stringency and for mapping reads originating from rRNA. Raw sequencing reads were first basecalled using bcl2fastq (version 2.20, Illumina). Sequencing adapters were removed and low quality read ends removed using trimmomatic^73^ with options ‘ILLUMINACLIP:adaptor.fa:4:30:10 LEADING:20 TRAILING:20 SLIDINGWINDOW:4:20 MINLEN:30’. A first round of PCR duplicate removal was performed using the RICpipe script ‘remove_PCR_duplicates.pl’ with default options. Forward and reverese reads where then mapped separately with STAR^74^ including the optimized mapping parameteres described in (REF CRSSN, options ‘--outFilterMultimapNmax 1 --outSAMattributes All --alignIntronMin 1 --scoreGap 0 --scoreGapNoncan 0 -- scoreGapGCAG 0 --scoreGapATAC 0 --chimSegmentMin 15 --chimJunctionOverhangMin 15 --limitOutSJcollapsed 10000000 --alignSJoverhangMin 15 --outSAMtype BAM SortedByCoordinate --chimOutType SeparateSAMold Junctions --outFilterScoreMinOverLread 0.66 --outFilterMatchNminOverLread 0.66 --chimFilter banGenomicN --chimScoreJunctionNonGTAG 0 --chimScoreDropMax 80 --chimNonchimScoreDropMin 20 --chimMainSegmentMultNmax 1 -- chimMultimapNmax 0 --alignIntronMax 24000000 --alignMatesGapMax 24000000’). Because STAR discriminates against chimeric reads located at different contigs, we generated a single contiguous reference genome that was used as mapping reference for STAR by concatenating all 14 nuclear chromosomes and masking all rRNA alleles except for one of the two identical *LSU_A_*(i.e. PF3D7_0725600 - PF3D7_0726000) and the *LSU_F_* rRNA locus (PF3D7_0801100 – PF3D7_0801200). Next, because STAR discriminates against backward chimeric alignments (i.e. on different strands), soft-clipped continuous alignments from the first round of STAR mapping were rearranged so that they can be re-mapped by STAR in a second round to increase the detection of backward chimeras. Soft-clipped reads were extracted using the CRSSANT script ‘softreverse.pl’ from the CRSSNT packages (https://github.com/zhipenglu/CRSSANT)^82^ using default settings, and the resulting forward and reverse reads were separately mapped again with STAR using the settings described above. Alignments derived from the first and second round of STAR alignment were processed in parallel from here on. Chimeric read pairs were next compiled from the separate forward and reverse alignment outputs using the ‘RICpipe’ script ‘collect_pair_tags.pl’, optical PCR duplicates were removed using samtools ‘fixmate’ and ‘rmdup’ and only read alignments ≥ 30nt and a maximum of one mismatch (SAM attribute ‘NM:i:0’ or ‘NM:i:1’) were retained. Chimeric alignments that were only detected in the second round of STAR alignment (i.e. those that were soft-clipped during the first round) were then combined with the filtered set of chimeric pairs already detected in the first round of STAR mapping, yielding a final set of read pairs per replicate. To add additional stringency to the mapping of reads to variant rRNA genes, we next extracted alignments overlapping *LSU_F_* and *LSU_A_*and mapped them again to the reference genome. Alignments for both loci were only retained if a read was reported to uniquely map to either one of them (STAR MAPQ value = 255). For the comparisons of RIC-seq derived interaction frequencies and spatial distances, (Fig. 2f), first the mean spatial coordinates of each carbon atom of a base in the 28S sequence were calculated from the A-type 80S ribosome model (PDB:3JBP). Pairwise distances for each coordinate were calculated in R and formatted to the cool format using juicer ‘pre’^83^ and hic2cool (https://github.com/4dn-dcic/hic2cool). All contact maps (Fig. 2f,h, Supplementary Fig. 3e) were visualized in R^77^ using package HiContacts^84^ with the resolution indicated in the legend, balanced scores and on a linear scale.

### rRNA fluorescent *in situ* hybridization and confocal imaging

Wild-type *P. falciparum* (strain NF54) gametes at 2 h.p.a. were obtained as described above. 1 mL of the culture was spun down and resuspended in freshly prepared 4% paraformaldehyde and 0.075% glutaraldehyde in DPBS, and incubated under slow rotation for 30 min. Fixed cells were washed 3 times in DPBS and stored at 4°C for a maximum of 2 weeks before staining and imaging. FISH probes for *LSU_F_* and *LSU_A_*rRNA were designed using the Stellaris Probe Designer (Biosearch Tenchnologies) following the approach describe in ref. ^85^. A set of 45 probes containing gene-specific targeting sequences as well as a shared FLAP-Y sequence were designed for *LSU_A_* rRNA, and a set of 40 probes containing a shared FLAP-X sequence were designed for *LSU_F_*rRNA (Supplementary Table 10). Primary probe sets were synthesized as 25 nmole DNA oligo plates (Integrated DNA Technologies). Secondary probes consisting of the corresponding FLAP-Y or FLAP-X antisense, conjugated with either Alexa Fluor 488 (FLAP-Y) or Alexa Fluor 594 (FLAP-X), were synthesized as 250 nmole DNA oligos in individual tubes (Integrated DNA Technologies). Primary gene-specific probes were combined in equimolar mixtures and pre-hybridized with the secondary fluorescent probes as described previously^85^. The hybridized duplexes were stored at -20°C until use. Cell permeabilization and staining was performed in accordance to the Stellaris RNA FISH protocol for cells in suspension (Biosearch Tenchnologies), beginning at the permeabilization step with 70% ethanol and using 3 μL of each hybridized duplex mix to prepare the Hybridization Buffer containing probe. Stained cells were resuspended in Vectashield Mounting Medium (Vector Laboratories no. H100010) and mounted on poly-D-lysine coated coverslips before imaging on a Zeiss LSM780 confocal microscope.

### ONT DRS rRNA modification detection

Gametes of *P. falciparum* wild-type parasites (strain NF54) were collected at 2 h.p.a. and total RNA was extracted as described. Total RNA samples were poly-A-tailed with E. coli Poly(A) polymerase (NEB no. M0276) and purified using RNAClean XP beads (Beckman Coulter no. A63987).

As a non-modified, negative control sample, we generated *in vitro* transcribed RNA of the *LSU_A_* and *LSU_F_*28S sequences. In brief, DNA was first extracted from asexual parasites using the DNeasy Blood and Tissue Kit (Qiagen no. 69504). Next, overlapping PCR fragments covering the 28S sequences of *LSU_A_* and *LSU_F_*were amplified using KAPA High Fidelity Polymerase (Kapa Biosystems no. KR0368) with the primers listed in Supplementary Table 10). Of note, in order to attach poly-A_(30)_ tails for subsequent *in vitro* transcription, poly-T_(30)_ tails were already included in the reverse primer of each PCR fragment. All PCR fragments were pooled at the same concentration in as input for a single *in vitro* transcription reaction using the MEGAscript Kit (Thermo Fisher no. AM1334). TURBO DNase provided with the kit was added to remove the eventual DNA template remains. Zymo Kit Clean and Concentrator kit (Zymo Research no. R1013) was used to purify the *in vitro* transcribed RNA before proceeding with libraries generation.

Libraries for the native, total RNA and IVT samples were prepared from ∼700 ng of input using the Direct RNA Sequencing Kit (ONT no. SQK-RNA004) following the manufacturer’s instructions and using the Induro reverse transcriptase (NEB no. M0681). Libraries were size seleted using RNAClean XP beads (Beckman Coulter no. A63987) in order to remove RNA fragments < 700 bp and the final libraries (∼900 ng) were sequenced using a FLO-MIN004RA flow cell (Nanopore Technologies) following the manufacturer guidelines on a Minion MK1 sequencer.

### ONT DRS read processing and alignment

Raw pod5 files were first merged using pod5 ‘merge’ (version 0.3.23, https://github.com/nanoporetech/pod5) and basecalled using dorado ‘basecaller’ (version 0.7.4, https://github.com/nanoporetech/dorado), with basecalling model ‘rna004_130bps_sup@v5.0.0 and options ‘--emit-moves --emit-sam’. Non primary and supplementary alignments were removed using samtools^75^ ‘view’ (option ‘-F 2304’), sorted using samtools ‘sort’, and only chromosome 7 and 8 (Pf3D7_07_v3 and Pf3D7_08_v3) that encode for the *LSU_A_* and *LSU_F_* rDNA were retained. The resulting BAM files were subsequently processed using the ‘align’ command of uncalled4^86^ (version 4.1.0). To get a visual overview of the current deviations from the model, heatmaps were generated using uncalled4 ‘trackplot’ with layer ‘dtw.model_diff’, targeting the peptidyl transferase center regions (Pf3D7_07_v3:1089773 – 1089880 and Pf3D7_08_v3:92036 – 91929). The normalized mean values of the signal current at a single-nucleotide resolution were obtained using uncalled4 ‘refstats’ with layer ‘dtw.current’. The differences in mean current between native samples (i.e. 2 h.p.a) and IVT control were plotted using ggplot2^79^ in R^77^ using the coordinates of the pairwise alignment in the abscissa (Fig. 2b). To calculate the histograms of current values for each position and read in the native and IVT control RNA (Fig. 2d), a tsv output file was generated using uncalled4 ‘align’ with option ‘--tsv-out’. The values corresponding to the positions of interest were extracted and visualized in R^77^ using ggplot2^79^. Alignments were visualized using the Integrative Genomics Viewer^72^.

To build a consensus rRNA modification dataset for *LSU_A_* and *LSU_F_*, first only the top 5% of sites (+/- 8 nt, the *k*-mer length read by the nanopore at any given time) with the highest difference between the native sample and the IVT control were retained. For *LSU_A,_* all of the 27 evolutionary conserved modifications detected by Pan-Mod-seq fall within this filter. For the corresponding positions (± 8nt) in *LSU_F_*, 10 did not fall within the top 5% of sites with the highest current difference. For the remaining 17, five of them that contain a base modification in *LSU_A_*were filtered out because the corresponding base at the alignment position in *LSU_F_* is different. This leads to a final set of 12 rRNA modifications in *LSU_F_* that 1) are within the top 5% of sites (± 8nt) with the largest difference in current between the native sample and IVT control 2) whose corresponding position in *LSU_A_* is also within the top 5% of sites (± 8nt) with the largest difference in current between the native sample and IVT control 3) whose corresponding position in *LSU_A_* is evolutionary conserved and 4) detected via Pan-Mod-seq.

### raPOOL

Three independent biological replicates were performed for the raPOOL experiments. Gametes of *P. falciparum* wild-type parasites (strain NF54) were enriched using NycoPrep and collected at 2 h.p.a. as described above. Per replicate, cell pellets of 2 mL of blood (1% parasitemia) were cross-linked as described for the RIC-seq analysis (see above) and snap-frozen in liquid nitrogen.

Parasite pellets were first resuspended in 500 μL of cold lysis buffer (50 mM tris-Cl, pH 7.0, 10 mM EDTA, and 1% SDS) and RNA was fragmented to ∼200 bp with a Bioruptor Pico sonication device (Diagenode no. B01080010, 15 cycles of 30 seconds ON/OFF, 4°C). Cell lysates were centrifuged (14,000 rcf for 10 min at 4°C) and the supernatant was kept for further processing. Affinity purification was carried out using the RNA affinity pool (‘raPOOL’) kit provided by siTOOLs. Here, the same probes that were designed for RNA-FISH targeting the 28S sequence of *LSU_F_* and *LSU_A_* (see above) were synthesized as pools of 3′-biotinylated single-stranded DNA oligos (siTOOLs). Following sonication and centrifugation, 1 mL of hybridization buffer was added to the cell lysate. In parallel, 15 μL of magnetic beads were prepared by washing trice with 1 mL of cell lysis buffer. To pre-clear the lysate and reduce non-specific binding, 15 μL of washed beads were added to 500 μL of cell lysate and incubated with rotation for 30 min at 37°C. Beads were then separated from the pre-cleared lysate using a magnetic rack. For hybridization, 2.5 μL of pooled raPOOL probes (100 pmol/µl) targeting either *LSU_F_* or *LSU_A_* were added to the pre-cleared lysate and incubated for 4 hours at 37°C with rotation. Next, 250 µl of washed beads were added and incubated with the cell lysate for 30 min at 37°C with rotation. Beads containing the RNA-protein complexes were separated on a magnetic rack and the supernatant was removed and stored for later processing. Beads were washed five times using 1 mL of pre-warmed wash buffer. During each wash, beads were gently agitated for 5 min and then separated on a magnetic rack. After the final wash, the beads were eluted in 95 μL proteinase K buffer (0.1M NaCl, 10mM TrisHCl pH 8.0, 1mM EDTA, 0.5% SDS). In parallel, 50 μL of the supernatant sample were mixed with 45 μL proteinase K buffer and processed in parallel. 5 μL of proteinase K were added to both samples and incubated for 45 min of incubation at 50°C with agitation to ensure complete protein digestion. The digested samples (i.e. supernatant and pulldown) were then mixed with TRIzol LS (Thermo Fisher no. 10296010) and RNA was extracted via isopropanol precipitation.

To remove the target *LSU_F_* or *LSU_A_*rRNA itself and further increase the coverage of mRNAs transcripts of interest, the pulldown samples were used as input for a second round of affinity purification. Here, the purified RNA extracted from the pulldown fraction was mixed with cold lysis buffer (final volume 500 μL), followed by hybridization with 1.5 μL of rRNA-targeting oligo pools and 150 μL of magnetic beads. Beads containing the target rRNA were separated on a magnetic rack, and RNA was extracted from the supernatant (i.e. rRNA depleted fraction) using TRIzol LS and isopropanol precipitation as described above.

Sequencing libraries were prepared using the NEBNext Ultra II Directional RNA Library Prep Kit (NEB no. E7760S), but omitting the initial fragmentation step. The final sequencing libraries were sequenced with a 2x150 bp paired-end layout on an Illumina NextSeq 2000 platform. Read pre-processing, alignment and filtering were performed as described above (mRNA sequencing analysis). Read counts were calculated using htseq-count^76^ using alignments that fall within exons (option ‘-t exon’). Differential abundance of mRNA transcripts in the two different conditions (i.e. supernatant and rRNA-depleted pulldown) were identified in R^77^ using package DESeq2^78^. *LSU_F_* target and repelled transcripts were filtered based on the adjusted *p*-value (*q* ≤ 0.05) and fold-change enrichment (rRNA-depleted pulldown over supernatant) ≥ 1.5 (i.e. *LSU_F_* ‘targets’) or ≤ -1.5 (i.e. *LSU_F_*‘repelled’). Coverage enrichments between the rRNA-depleted pulldown and supernatant samples were calculated using deeptools^87^ ‘bamCompare’ and metagene analyses (Fig. 3g, Supplementary Fig. 4i,j) were performed in R^77^ using package ‘chipseeker’^88^.

---

**Supplementary Figure 1.**
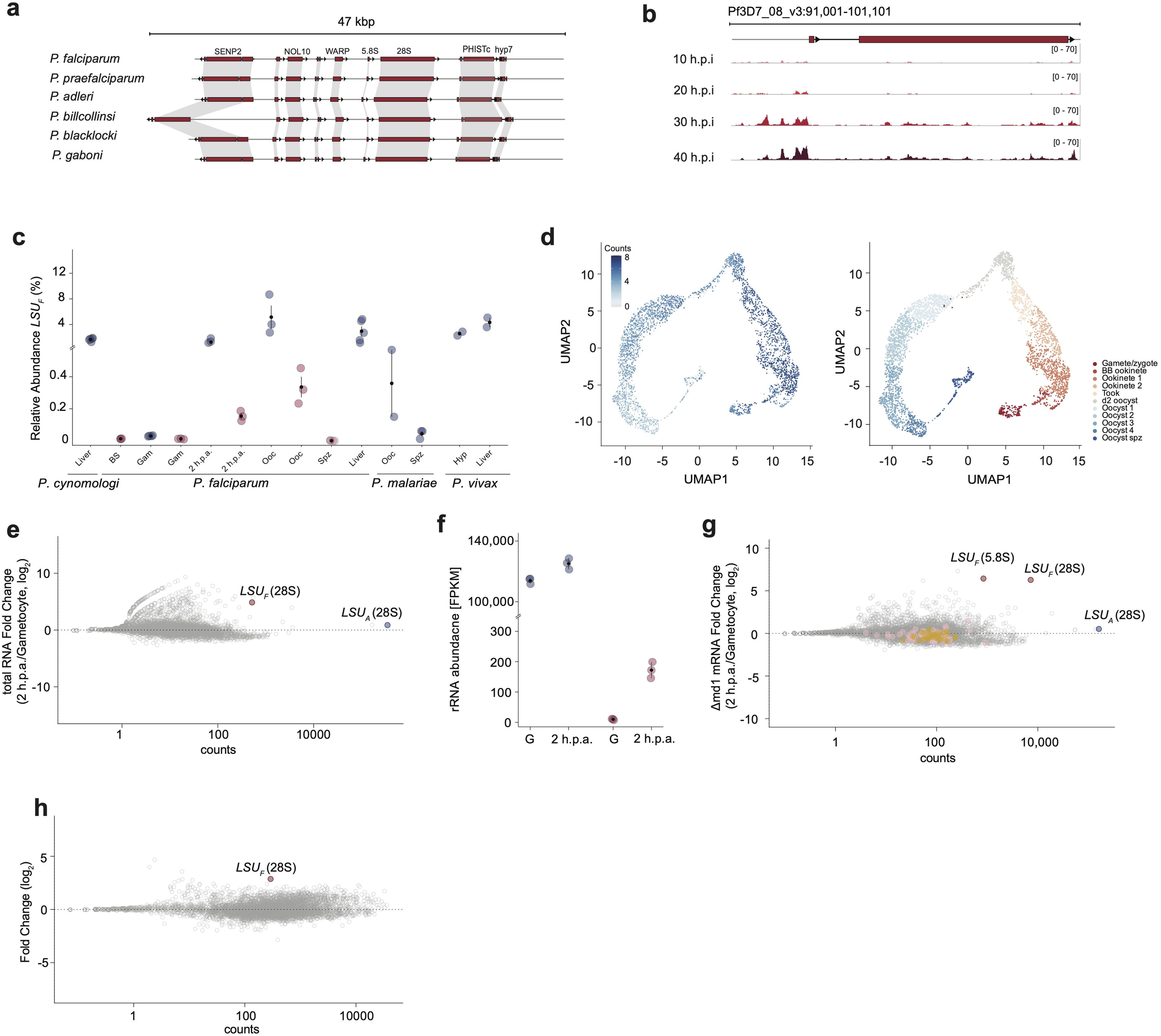
(a) Synteny of the *LSU_F_* rDNA locus showing a conserved genomic positioning across species of the *Laverania* class. (b) ATAC-seq (assay for transposase-accessible chromatin with sequencing) ratio tracks (ATAC/gDNA input)^55^ across the asexual replicative lifecycle of *P. falciparum*. h.p.i.: Hours post invasion. (c) Relative rRNA expression levels of *LSU_F_* homologs (calculated as *LSU_F_* [FPKM] / other rRNA [FPKM]) in *P. falciparum*^12,56,57^*, P. malaria*^58^*, P. cynomologi*^59^ and *P. vivax*^60^ measured in different stages of parasite development. Expression levels measured via mRNA- or total RNA-seq are highlighted in blue and red, respectively. (d) scRNA-seq expression profile of *P. falciparum LSU_F_* 28S (PF3D7_0726000) during development in the *An. gambiae* mosquito (left) and annotation of stages during *P. falciparum* development in the *An. gambiae* mosquito^20,25^ (right). (e) Changes in transcript abundance in wild-type parasites between the gametocyte stage and 2 h.p.a. calculated from total RNA sequencing. *LSU_A_* and *LSU_F_* 28S transcripts are shown in red and blue, respectively. (f) Comparison of absolute rRNA abundance (measured as fragments per kilobase of exon per one million mapped reads, FPKM; calculated from total RNA-seq) for *LSU_A_* and *LSU_F_*28S rRNA in wild-type gametocytes (G) and gametes at 2 h.p.a. (g) Changes of transcript abundance between Δmd1 gametocytes and gametes at 2 h.p.a. calculated from mRNA-seq. Annotated large and small subunit ribosomal proteins are highlighted in pink and yellow, respectively. (h) Changes in transcript abundance in wild-type parasites (strain NF54) between the gametocyte stage and 5 min post activation (m.p.a.) calculated from mRNA sequencing. *LSU_A_* and *LSU_F_* 28S are shown in red and blue, respectively^30^.

**Supplementary Figure 2.**
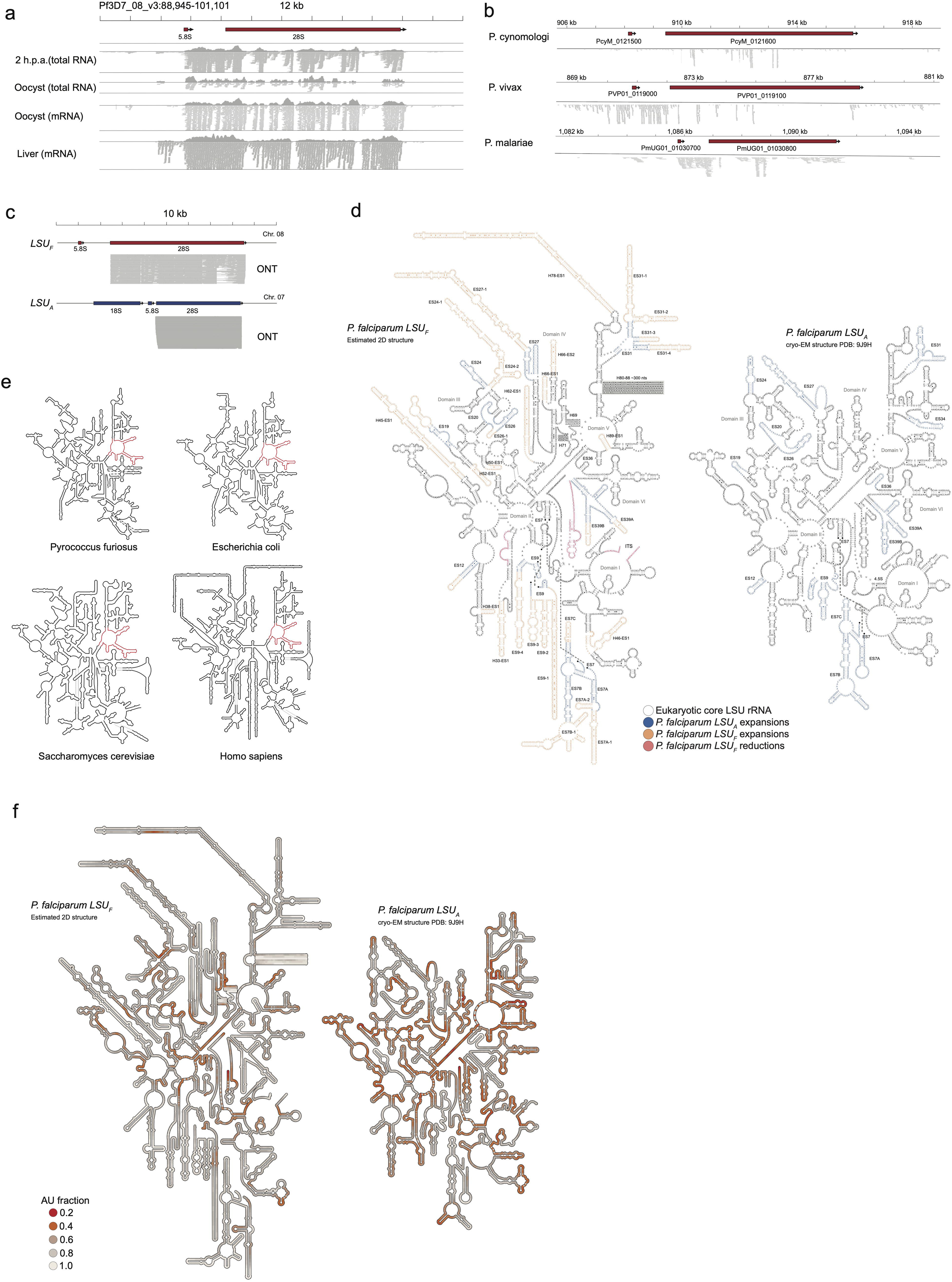
(a) Alignments and read coverage across *LSU_F_* rRNA at the indicated developmental stages derived from short-read mRNA and total RNA-seq experiments.^12,56,57^ (b) Alignments and mRNA read coverage across *LSU_F_* homologs showing the retention of the ITS2 sequence. All data are derived from parasites during hepatocyte development.^58–60^ (c) Alignments and coverage of direct RNA sequencing reads of the *in vitro* transcribed control RNA fragments used as negative (i.e. non-modified) control for *LSU_F_*(top) and *LSU_A_* (bottom). (d) Secondary structure map of *LSU_F_*(left) and the canonical *LSU_A_* (right) rRNA including nucleotide identities and numbering of expansion segments (ES). H: Helix (e) Exemplary secondary structure maps of LSU rRNA from archeae (*Pyrococcus furiosus*), bacteria (*Escherichia coli*), fungi (*Saccharomyces cerevisiae*) and metazoans (*Homo sapiens*)^61,62^. The peptidyl transferase center is highlighted in red. (f) Secondary structure map of *LSU_A_*and *LSU_F_* rRNA colored by AU content.

**Supplementary Figure 3.**
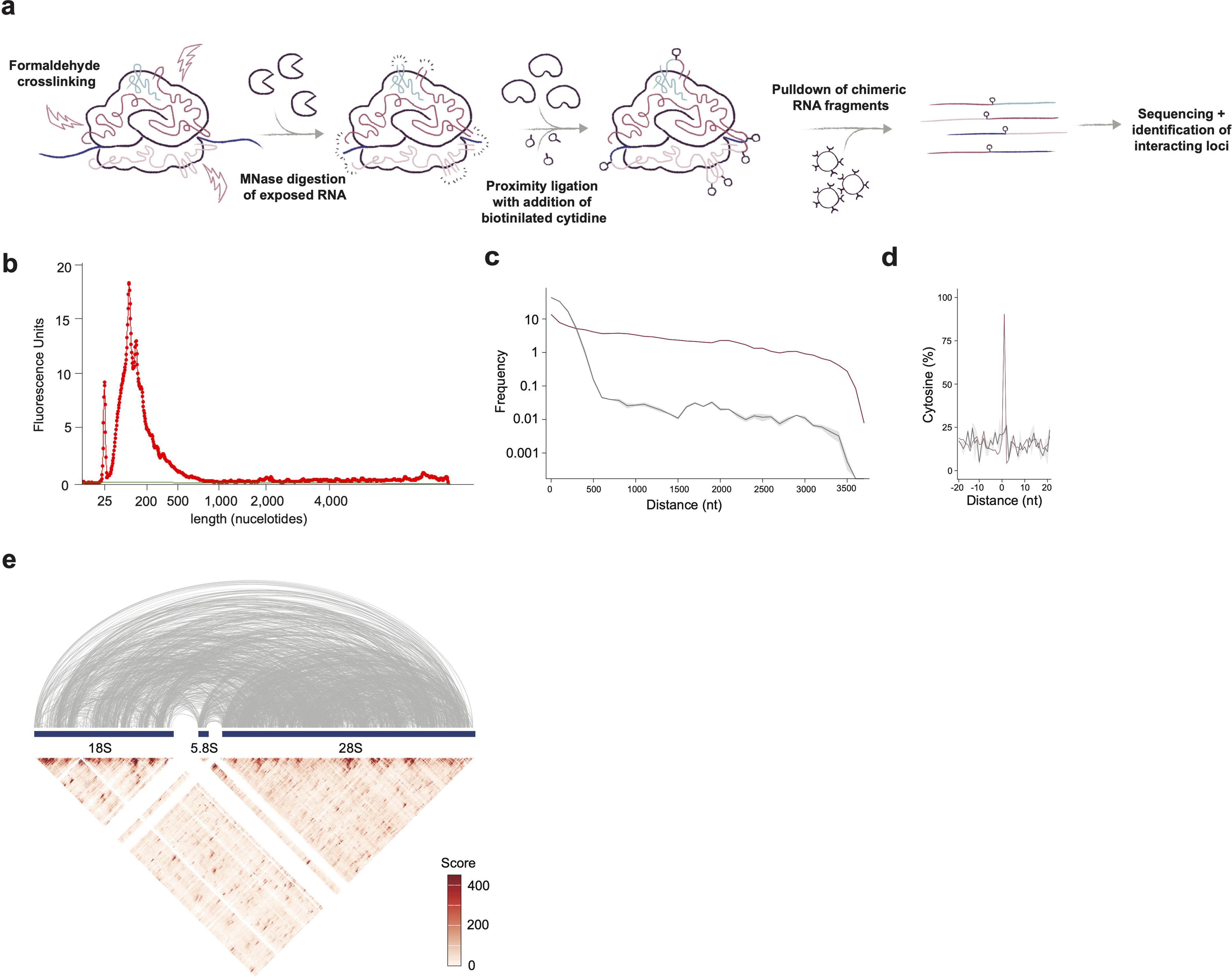
(a) Workflow of the RIC-seq experiment to detect intra- and intermolecular interactions of *LSU_F_*and *LSU_A_* rRNA. (b) Fragment length of cross-linked RNA after MNase digestion and prior to proximity ligation. (c) Intramolecular contact frequency by distance (nucleotides) within the *LSU_A_* 28S of the RIC-seq (red) or control samples (grey). Long-range interactions are frequently detected by RIC-seq, whereas the control sample predominantly recovers interactions of neighboring regions. (d) Percentage of cytosines detected at the junction of chimeric reads in the merged RIC-seq dataset (red) or the merged control sample dataset (grey). (e) Intra- and Intermolecular interactions within and between the *LSU_A_* 18S, 5.8S and 28S sequences (top). Balanced contact frequencies of intra- and intermolecular interactions among the different A-type rRNA detected by RIC-seq. Linear scale, resolution: 5 nt.

**Supplementary Figure 4.**
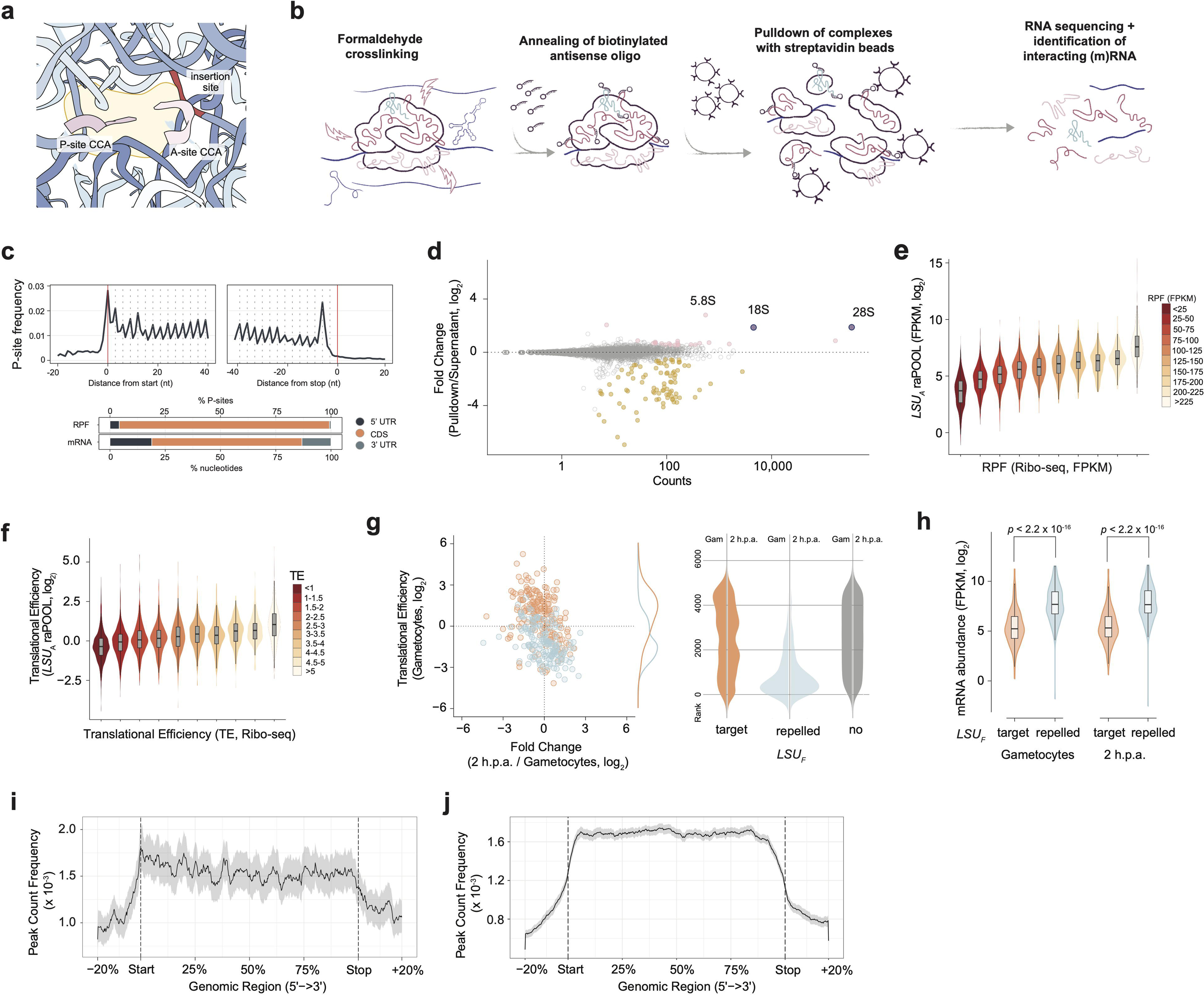
(a) View into the ribosomal exit tunnel of the canonical *P. falciparum* A-type ribosome (PDB: 9J9H). The insertion site of the *LSU_F_* PTC hairpin is highlighted in red, the 3’ CCA ends of the tRNAs located at the A- and P-site are highlighted in pink (see also Fig. 3c). (b) Workflow of the raPOOL experiment to enrich for mRNA transcripts bound to *LSU_F_* or *LSU_A_*rRNA. (c) Top: Meta-profile showing the trinucleotide periodicity of ribosomes along the first and last 40 nt of coding sequences on a transcriptome-wide level in wild-type (strain NF54) parasites sampled at 2 h.p.a. Bottom: Relative distribution of P-sites in the 5’UTR, 3’UTR and CDS across the transcriptome in wild-type (strain NF54) parasites sampled at 2 h.p.a. The cumulative length of each feature across the transcriptome is shown on the right, indicating a strong bias for P-site identification within the CDS. (d) Relative enrichment and depletion of mRNAs associated with the *LSU_A_* 28S rRNA (blue). mRNAs that are significantly enriched (*q* ≤ 0.05. fold-change ≥ 1.5) or depleted (*q* ≤ 0.05. fold-change ≤ -1.5) in the *LSU_A_* raPOOL experiment are highlighted in pink and yellow, respectively. (e) Correlation between normalized read coverage of *LSU_A_*-associated mRNA identified by raPOOL (ordinate) and coverage of ribosome protected fragments (RPFs) identified by Ribo-seq (abscissa). (f) Correlation between translation efficiencies calculated from *LSU_A_*-associated mRNA identified by raPOOL (*LSU_A_* raPOOL_FPKM_ / mRNA_FPKM_: ordinate) and translation efficiencies calculated by Ribo-seq (RPF_FPKM_ / mRNA_FPKM_, abscissa). (g) Left: Comparison of translation efficiency in gametocytes (ordinate) and change thereof during parasite transmission (abscissa) for *LSU_F_* target (orange) and repelled transcript (blue). Right: Distribution of ranked absolute translational efficiencies in gametocytes (left) and gametes at 2 h.p.a. (right) for *LSU_F_*targets (orange), repelled (blue) and other transcripts (grey). (h) Comparison of transcript abundance (measured as FPKM) of *LSU_F_* target and repelled mRNAs in mature gametocytes (left) and at 2 h.p.a. (right). *p*-values were calculated with an unpaired Wilcoxon Rank Sum test. (i) Metagene plot showing the average enrichment of *LSU_A_* associated mRNA reads (pulldown/supernatant) across mRNA transcripts. Grey area indicates 95% confidence interval. (j) Metagene plot of average RPF coverage across mRNA transcripts at 2 h.p.a. for wild-type (strain NF54, left) parasites. Grey area indicates 95% confidence interval.

**Supplementary Figure 5.**
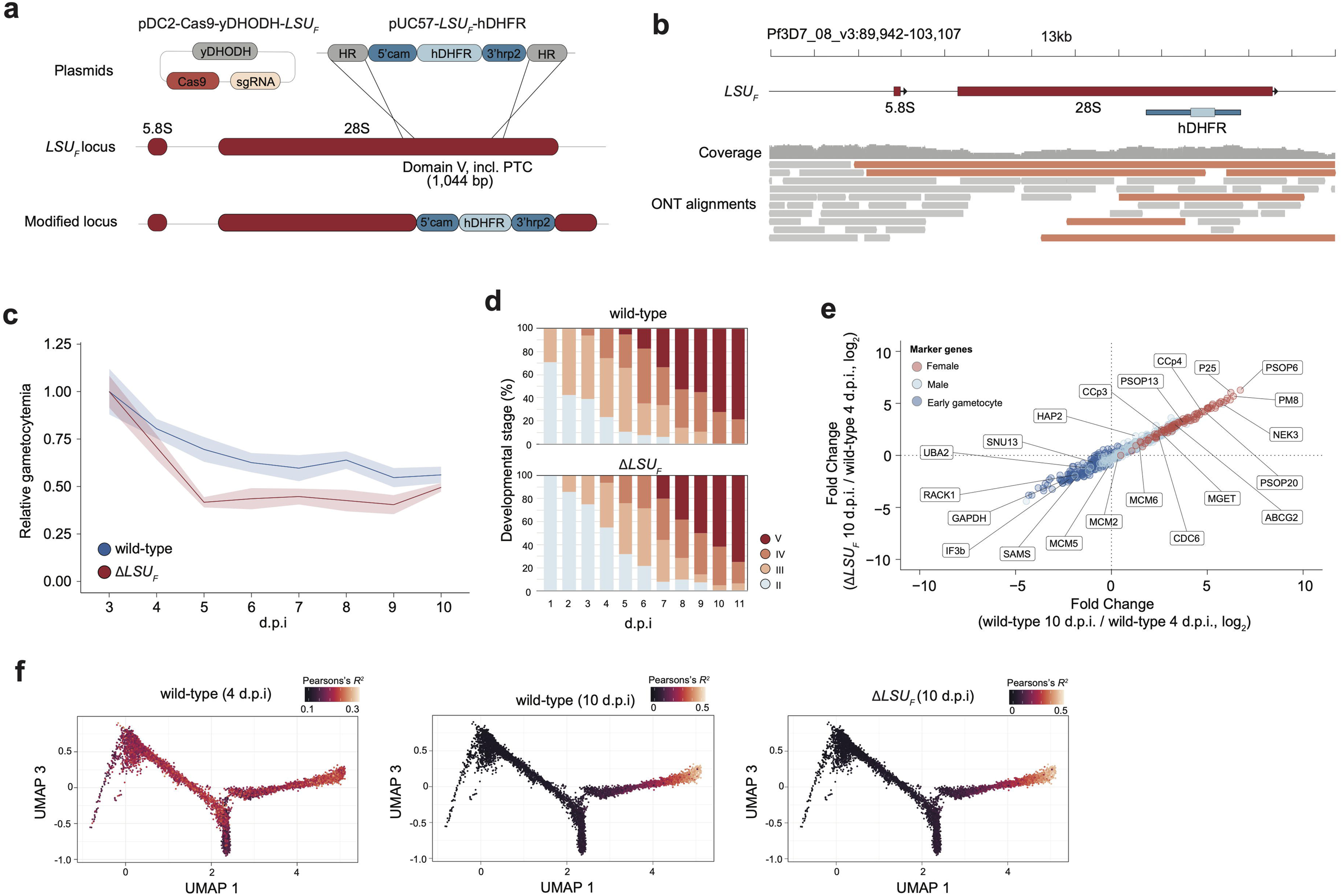
(a) Representation of the strategy to remove 1,044 bp of the *LSU_F_* 28S rRNA (PF3D7_0801100: 4,389 – 5,433) and replacement with a hDHFR resistance cassette. 5’ cam: Calmodulin promoter; hDHFR: human dihydrofolate reductase; 3’ hrp2: histidine-rich protein 2 3’ untranslated region; HR: Homology region; yDHODH: yeast dihydroorotate dehydrogenase. (b) Nanopore tracks displaying read alignments encompassing the modified *LSU_F_* rDNA locus (highlighted in orange), showing successful genome editing. (c) Proportion of gametocyte parasitemia (DNA stained with DAPI) relative to day 3 post gametocyte induction as measured by flow cytometry; asexual parasites were absent starting from day 4 post gametocyte induction. Data are presented as mean (bold lines) +/− 1 standard deviation (ribbon) calculated from 3 biological replicates. d.p.i.: days post induction (d) Relative abundance of developmental stages during gametocytogenesis for wild-type (strain NF54, top) and Δ*LSU_F_* (bottom) parasites showing no phenotypic differences when *LSU_F_* is deleted. d.p.i.: days post induction (e) Comparison of marker gene expression changes of immature wild-type (strain NF54) parasites (sampled at day 4 post gametocyte induction) to mature wild-type (strain NF54, abscissa) and Δ*LSU_F_* (ordinate) parasites. Mature gametocytes were sampled at day 10 post gametocyte induction. Immature (blue), male (turquoise) and female (red) marker genes were taken from ref. ^63^ (f) Developmental age of wild-type (strain NF54) and Δ*LSU_F_* parasites derived from the correlation of mRNA-seq and a scRNA-seq reference dataset^63^. Each dot represents a scRNA-seq transcriptome along gametocytogenesis, with the color indicating the Pearson’s *R*^2^ correlation co-efficient between the transcriptomes wild-type/Δ*LSU_F_* transcriptomes and the individual scRNA-seq transcriptome.

**Supplementary Figure 6.**
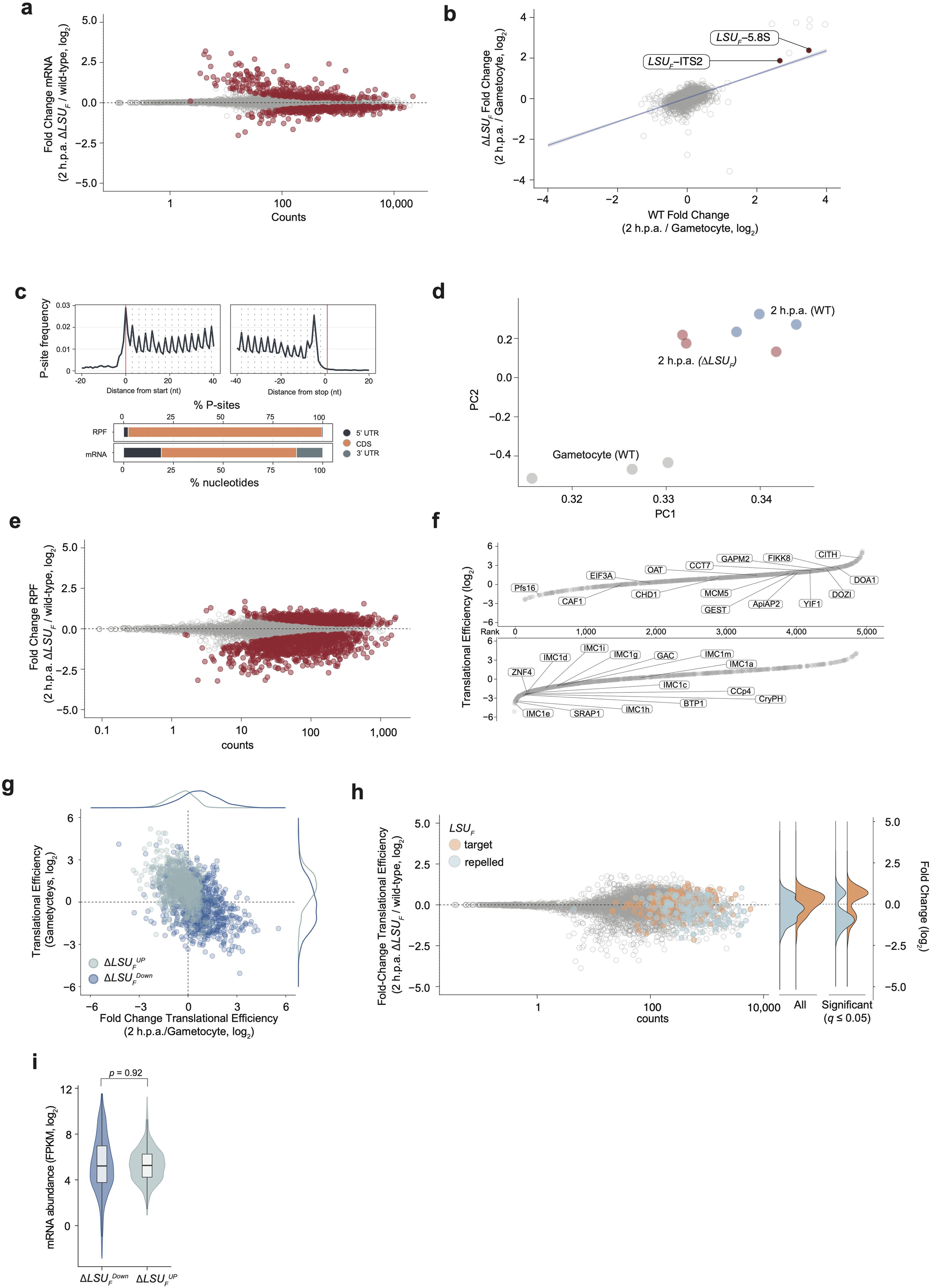
(a) Changes in mRNA abundance between wild-type (strain NF54) and Δ*LSU_F_* parasites at 2 h.p.a. Transcripts featuring significant changes in mRNA abundance (*q* ≤ 0.05) are shown in red. (b) Comparison of changes in mRNA abundance (represented as log_2_ fold changes) between gameto cytes and gametes at 2 h.p.a in wild-type (strain NF54, abscissa) and Δ*LSU_F_* (ordinate) parasites showing similar transcriptional dynamics and upregulation of *LSU_F_* rRNA regions located upstream of the removed *LSU_F_* region in Δ*LSU_F_* (i.e. *LSU_F_* 5.8S and ITS2 rRNA, red). (c) Top: Meta-profile showing the trinucleotide periodicity of ribosomes along the first and last 40 nt of coding sequences on a transcriptome-wide level in Δ*LSU_F_* parasites sampled at 2 h.p.a. Bottom: Relative distribution of P sites in the 5’UTR, 3’UTR and CDS across the transcriptome in Δ*LSU_F_* parasites sampled at 2 h.p.a. The cumulative length of each feature across the transcriptome is shown on the right, indicating a strong bias for P-site identification within the CDS. (d) PCA plot calculated from translation efficiencies in mature wild-type gametocytes and wild-type (strain NF54, left) and Δ*LSU_F_* (right) parasites 2 h.p.a indicating successful re-initation of development and transition of the general translational program. (e) Changes in ribosome protected footprint abundance between wild-type (strain NF54) and Δ*LSU_F_* parasites at 2 h.p.a. Transcripts featuring significant changes in ribosome protected footprint abundance are shown in red. (f) Ranking (abscissa) of translational efficiencies in gametocytes (ordinate) for mRNA transcripts with significantly increased (Δ*LSU_F_*^UP^, top) or decreased (Δ*LSU_F_*^DOWN^, bottom) rates of translation at 2 h.p.a. following *LSU_F_*deletion. See also Fig. 4b (g) Comparison of translation efficiencies in gametocytes (ordinate) and change thereof during parasite transmission (abscissa) for Δ*LSU_F_*^UP^ (turquoise) and Δ*LSU_F_*^DOWN^ (blue). (h) Changes in translational efficiencies between wild-type and Δ*LSU_F_* parasites at 2 h.p.a. as detected by Ribo-seq (see also Fig. 4a). *LSU_F_* target and *LSU_F_* repelled mRNA transcripts are highlighted in orange and turquoise, respectively. Right: Distribution of log_2_ fold changes of translational efficiency for all *LSU_F_* target and repelled mRNA transcripts (left) and those featuring significant changes in translational efficiency (right), indicating that *LSU_F_* targets more often feature higher translational efficiencies when *LSU_F_* is depleted and *vice versa* for *LSU_F_* repelled mRNA transcripts. (i) Transcript abundance (in FPKM) of Δ*LSU_F_*^DOWN^ (blue, left) and Δ*LSU_F_*^UP^ transcripts (turquoise, right) in wild-type (strain NF54) gametocytes, indicating overall similar abundance of the two mRNA classes in the gametocyte stage. *p-*value was calculated using an unpaired Wilcoxon Rank Sum test.

